# L-type channel voltage-dependent facilitation results from asymmetric π-H and π-π quadrangle interactions at DI–DII domains

**DOI:** 10.64898/2026.01.23.701029

**Authors:** Aynal Hoque, Vishal Tanaji Khade, Venkata Gopal Ede, Abhijeet S. Kate, Gerald W Zamponi, Giriraj Sahu

**Affiliations:** Neuronal Excitability and Disease Research Group, Molecular Biophysics Unit, Indian Institute of Science, Bengaluru, India; Department of Biotechnology, National Institute of Pharmaceutical Education and Research, Ahmedabad, Palaj, Gandhinagar, India; Department of Natural Products, National Institute of Pharmaceutical Education and Research, Ahmedabad, Palaj, Gandhinagar, India; Department of Clinical Neurosciences, Hotchkiss Brain Institute and Alberta Children’s Hospital Research Institute, University of Calgary, Calgary, Canada

**Keywords:** Voltage-gated calcium channels, L-type channels, Voltage-dependent facilitation (VDF), Patch clamp recording, Single-channel recordings, Molecular dynamics, Leachables, 2, 4-DTBP

## Abstract

Voltage-dependent facilitation (VDF) is a distinctive phenomenon of L-type voltage-gated calcium (Ca_V_) channels, in which depolarized prepulse voltages increase inward current by up to several fold. Despite extensive study, the molecular origin and biophysical mechanisms underlying the L-type channel (Ca_V_1.1-Ca_V_1.4) VDF remain elusive. A serendipitous observation that solutions stored in polypropylene tubes eliminated Ca_V_1.2 and Ca_V_1.3 L-type channel VDF led us to identify the leachable 2,4-di-tert-butylphenol (2,4-DTBP) as a selective VDF inhibitor. Docking and molecular dynamics simulations revealed previously unreported asymmetric π-H and π-π interdomain quadrangle interactions at the DI–DII pore-domain (PD) interface in L-type channels. Disruption of these quadrangle interactions in Ca_V_1.2 via point mutagenesis abolished the VDF. Whereas, introducing the quadrangle interaction onto non-VDF-displaying Ca_V_2.1 channels induced robust VDF. This work provides the first molecular evidence for the VDF endpoint: L-type channel-specific interdomain quadrangle interactions asymmetrically positioned at the DI–DII PD interface.

## Introduction

Voltage-gated calcium (Ca_V_) channels are multisubunit proteins that play crucial physiological roles in regulating membrane depolarization and transmission of action potentials in brain neurons, cardiomyocytes, muscle, and endocrine cells^1–3^. The primary pore-forming α_1_subunit in Ca_V_ channels consists of four domains (D1–DIV), with each bearing six transmembrane segments (S1–S6). S1–S4 constitute the voltage-sensing domain (VSD), and S5–S6 form the pore domain (PD), with the P-loop, flanked by two short α helices (P1 and P2)^4–7^. Physiologically, the functions of Ca_V_ channels are fine-tuned by various activity-dependent regulatory mechanisms that determine the magnitude and kinetics of intracellular Ca^2+^ entry. Alterations in Ca_V_ channel function have been linked to cardiovascular, neuropsychiatric, and neurodegenerative diseases^1,8–11^.

Among the high-voltage-activated (HVA) Ca_V_ channels (comprising Ca_V_1.1-1.4 and Ca_V_2.1-2.3)^1,2,12^, L-type channels (Ca_V_1.1-1.4) uniquely exhibit activity-induced depolarizing prepulse (DPP) mediated voltage-dependent facilitation (VDF) of intracellular Ca^2+^ entry, a phenomenon that is not observed in Ca_V_2 channels^13–17^. In contrast to VDF, both Ca_V_1 and Ca_V_2 channels exhibit other forms of facilitation, such as calcium-dependent facilitation (CDF) and RGK-GTPase-mediated facilitation^18–24^. The magnitude of VDF is calcium-independent but graded with the amplitude of the prepulse voltages, where a DPP of varying magnitude, duration, and frequency shifts the gating transitions to a state of long-duration higher open probability^13,15–17,25,26^. L-type channel VDF facilitates inward Ca_V_1 currents during repeated action potentials in cortical, striatal, cerebellar, and dorsal root ganglion neurons and contributes to secretion of catecholamines, exocytosis, intensification of muscle contraction, and the automaticity of sino-atrial myocytes during cardiac autorhythmicity^15,16,27–31^. Despite being of critical physiological importance, the mechanisms underlying the L-type Ca_v_1 channel VDF remain unanswered. Expression systems using the Ca_V_1.2 and Ca_V_1.3 cDNAs have attempted to delineate the biophysical mechanisms underlying the VDF, but with limited success. Chimeric constructs comprising Ca_V_1.2 domains with non-VDF-displaying Ca_V_2.3/Ca_V_2.1 have demonstrated the dominant role of DI and DII in attaining VDF^14,32^. However, the molecular endpoints and biophysical processes underlying DPP-induced VDF in L-type channels remain elusive. Furthermore, the molecular mechanisms that enable L-type channels to uniquely display VDF, but not other HVA channels, remain to be fully elucidated.

In this work, serendipitously, we observed that buffer solutions stored in laboratory polypropylene (PP) plastic Falcon tubes markedly eliminated the DPP-stimulated VDF of Ca_V_1.2 and Ca_V_1.3 L-type channels, leaving the activation and inactivation kinetics intact. These observations, however, provided an opportunity to unravel the molecular endpoint contributing to the mysterious L-type channel VDF. Leachable characterization, followed by molecular dynamics (MD) simulations and electrophysiology, revealed potent VDF inhibition of the Ca_V_1.2 and Ca_V_1.3 L-type channels by the plastic leachable 2,4-di-tert-butylphenol (2,4-DTBP). The inhibition of VDF was achieved by disrupting a previously unreported Ca_V_1 channel-specific, crucial asymmetric π-H and π-π quadrangle interaction at the DI–DII PD interface. Whole-cell and single-channel recordings in the alanine-substituted Ca_V_1.2 mutated channels demonstrated the absolute requirement of the quadrangle interactions in producing VDF. Notably, incorporation of π-H and π-π quadrangle interactions into non-VDF facilitative Ca_V_2.1 channels transformed them to produce robust VDF. Collectively, these findings discovered and characterized a novel asymmetric π-H and π-π quadrangle interdomain interaction that underlies the molecular endpoint of activity-induced VDF, a phenomenon unique to L-type HVA channels.

## Results

### L-type channel VDF is inhibited by PP stored solutions

To gain deeper insight into the molecular endpoints of L-type channel VDF, we coexpressed the primary α subunit of either Ca_V_1.2 (CACNA1C, α_1C_) or Ca_V_1.3 (CACNA1D, α_1D_) in tsA-201 cells along with the accessory subunit β_1b_(CACNB1) in a 1:0.75 ratio. In a separate set of experiments, α_1C_ or α_1D_ cDNAs were coexpressed with β_1b_, along with the α_2_δ_1_ (CACNA2D1) accessory subunit in a 1:0.75:0.5 ratio. Whole-cell patch-clamp recordings were performed on cells held at -70 mV and bathed in an external solution containing 5 mM BaCl_2_, stored in glass reagent bottles. Voltage steps ranging from -80 mV to +60 mV with 10 mV increments resulted in typical slowly inactivating inward Ca_V_1.2 and Ca_V_1.3 currents with a half maximal activation voltage (V_a_) of -7.13 ± 1.98 mV (n = 9) and -26.63 ± 3.65 mV (n = 8), respectively, in cells coexpressing β_1b_ (***Fig. S1A-S1D***), and V_a_ of -13.50 ± 0.64 mV (n = 9) and - 27.59 ± 1.20 mV (n = 6), respectively, in cells coexpressing α_2_δ_1_and β_1b_ (***Fig. S2A-S2D, Table S1***)^33^. For eliciting the VDF of Ca_V_1.2 and Ca_V_1.3 channels, a series of DPP voltages ranging from 0 to 180 mV was applied preceding a test pulse (P2) and compared with a non-pairing pulse (P1)^13,14,34,35^. The magnitude of VDF was measured by the difference between the peak current amplitudes of P2 and P1 stimuli. Both Ca_V_1.2 (***Fig. 1A and 1C***) and Ca_V_1.3 (***Fig. 1B and 1D***) coexpressed with β_1b_, exhibited a substantial increase in P2 current magnitudes in response to 0-180 mV DPP stimulus, reaching a maximum of ∼65% and ∼45%, respectively, at DPP of 100-120 mV in external solutions stored in glass reagent bottles. The results are consistent with the reported DPP-induced facilitation of Ca_V_1.2^13–15,34,36^ and Ca_V_1.3 channels^29,31,37^. Moreover, coexpression of α_2_δ_1_ along with β_1b_ subunit resulted in a comparatively lower VDF in Ca_V_1.2 (∼36% at 120 mV, p = 0.0054) (***Fig. S3A, S3C, and S3E***) but no difference was observed in Ca_V_1.3 (∼42% at 120 mV, p = 0.82) (***Fig. S3B, S3D, and S3E***) (***Table S1***). In earlier reports, the inclusion of α_2_δ_1_was reported to inhibit the VDF of Ca_V_1.2^38^. The observed deviation in this work might be due to the 1:0.5 α_2_δ_1_ cDNA used in relation to Ca_V_1.2, compared with a 1:1 ratio used in the earlier study. Indeed, an increase in α_2_δ_1_ from a 1:0.5 to a 1:1 ratio in relation to Ca_V_1.2 significantly decreased the VDF (***Fig. S4***), suggesting that the VDF of Ca_V_1.2 channels is dependent on the degree of α_2_δ_1_engagement. α_2_δ_1_also markedly increased the current inactivation, apparent from a decreased time constant of inactivation (τ, msec), and residual current plot (ratio of peak current to current at 400 msec, R_400_) of Ca_v_1.2 (***Fig. S1E and S2E)*** and Ca_V_1.3 ***(S1F and S2F***) (***Table S1***), in agreement with earlier observations^37,39^.

**Figure 1:**
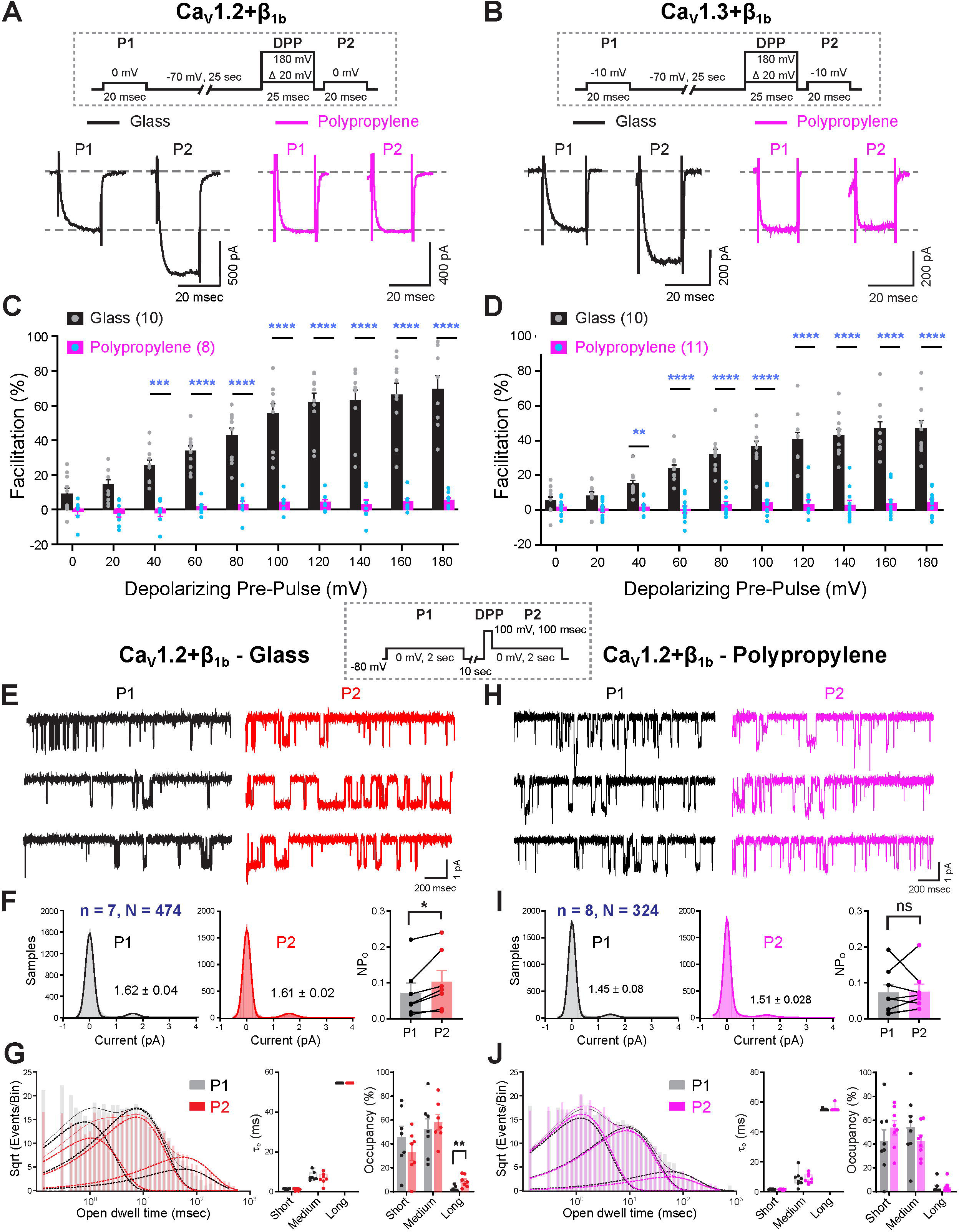
Buffer solutions stored in PP tubes inhibit the VDF of L-type channels: **(A and B)** Representative whole-cell current traces of L-type Ca_V_1.2 **(A)** and Ca_V_1.3 **(B)** channels coexpressed with β_1b_ subunit in tsA-201 cells recorded in external solutions stored in glass bottles (black) or polypropylene (PP) tubes (magenta). The voltage protocol is shown in the dotted box above. **(C and D)** Bar plot quantifications depict percentage facilitation of Ca_V_1.2 **(C)** and Ca_V_1.3 **(D)** whole-cell currents, measured by the difference between peak current amplitudes of P2 and P1 (P2-P1) voltage steps, in buffer solutions stored either in glass bottles (black) or PP tubes (magenta). Experimental replicate cell numbers are indicated in brackets. Two-way ANOVA is used for statistical significance. **(E and H)** Exemplary cell-attached single-channel current traces of Ca_V_1.2 coexpressed with the accessory subunit β_1b_ in tsA-201 cells recorded in external solutions stored in glass bottles **(E)** or PP tubes **(F)**. The voltage protocol used is shown in the dotted box above. **(F and I)** The histogram plots show the single-channel current amplitudes, while the bar plots display the open probability of Ca_V_1.2 cell-attached recordings obtained at 0 mV step of 2 sec in buffer solutions stored in glass bottles **(F)** or PP tubes **(I)** before (P1) or after (P2) the DPP to 100 mV for 100 msec. n = number of cells, N = number of single-channel traces. A non-parametric paired Student’s t-test is used to determine statistical significance. **(G and J)** The logarithmic histogram plots show the open dwell-time distributions of Ca_V_1.2 channels in glass or PP-stored solutions, fitted by nonlinear least squares, while the bar plots display open time constants and their occupancy in short, medium, and long open states. One-way ANOVA with pairwise comparison was used to determine statistical significance. All bar plots represent mean ± SEM. *p ≤ 0.05, **p ≤ 0.01, ***p ≤ 0.001, ****p < 0.0001, ns > 0.05; ns, non-significant.

Serendipitously, we observed a near blockade in the potentiation of the P2 current amplitudes of Ca_V_1.2 and Ca_V_1.3 upon stimulation with 0-180 mV DPP in cells coexpressing β_1b_ (***Fig. 1A-1D***) or β_1b_and α_2_δ_1_(***Fig. S3A-S3D***) in external solutions prepared with 1M BaCl_2_ stock that was stored in laboratory PP tubes for 1-2 weeks. The maximum VDF for these channels was 3-5%, indicating nearly complete inhibition of this process (***Table S1***). Lack of VDF in Ca_V_1.2 and Ca_V_1.3 channels can be observed as early as 5-7 days after storage of BaCl_2_ stock solution in PP tubes or external solutions prepared from Double-distilled (DD) water stored in PP tubes, indicating an inhibition of VDF plausibly due to plastic leachables^40–42^. Further to characterize whether PP-stored solutions indeed inhibited the VDF of L-type channels or rendered the channels in a prefacilitated state, Ca_V_1.2 VDF was examined in response to +140 mV DPP in glass bottle stored solutions, followed by 5-10 min incubation in PP-stored solutions on the same cells, resulting in a complete disappearance of VDF without affecting the basal current density (***Fig. S5A and S5B***), demonstrating an inhibition of VDF by PP-stored solutions rather than transforming the channels to a prefacilitated state. Moreover, no noticeable change was observed in the current density, voltage dependence, inactivation kinetics, and residual R_400_ current pattern of Ca_V_1.2 and Ca_V_1.3 channels coexpressed with β_1b_(***Fig. S1A-S1F***) or β_1b_ and α_2_δ_1_ (***Fig. S2A-S2F***), recorded in external solutions stored in PP tubes (***Table S1***). These results highlight a specific effect of plastic leachables on blocking the VDF of L-type Ca_V_1.2 and Ca_V_1.3 channels, with all other gating properties remaining intact.

### PP stored solutions inhibit the DPP-induced gating shift

Along with an increase in current density, earlier examinations reported a hyperpolarizing shift in the Ca_V_1.2 activation curve due to DPP stimulation^13,16,43^. To validate the inhibition of L-type channel VDF by PP-stored solutions, we examined Ca_V_1.2 activation from -80 to +60 mV, with or without a DPP of +140 mV in cells coexpressing β_1b_. Control cells bathed in glass bottle stored solutions showed a significant hyperpolarizing shift in the Ca_V_1.2 I-V plot that reflected in a ∼-5 mV negative shift in V_a_, which is completely inhibited in cells bathed in PP incubated solutions (***Fig. 2A***). Also no significant change in V_a_ of Ca_V_1.2 was observed between glass and PP stored solutions without DPP stimulus, indicating an inhibition of VDF rather than modifying the basal activation or attaining a prefacilitated state.

**Figure 2:**
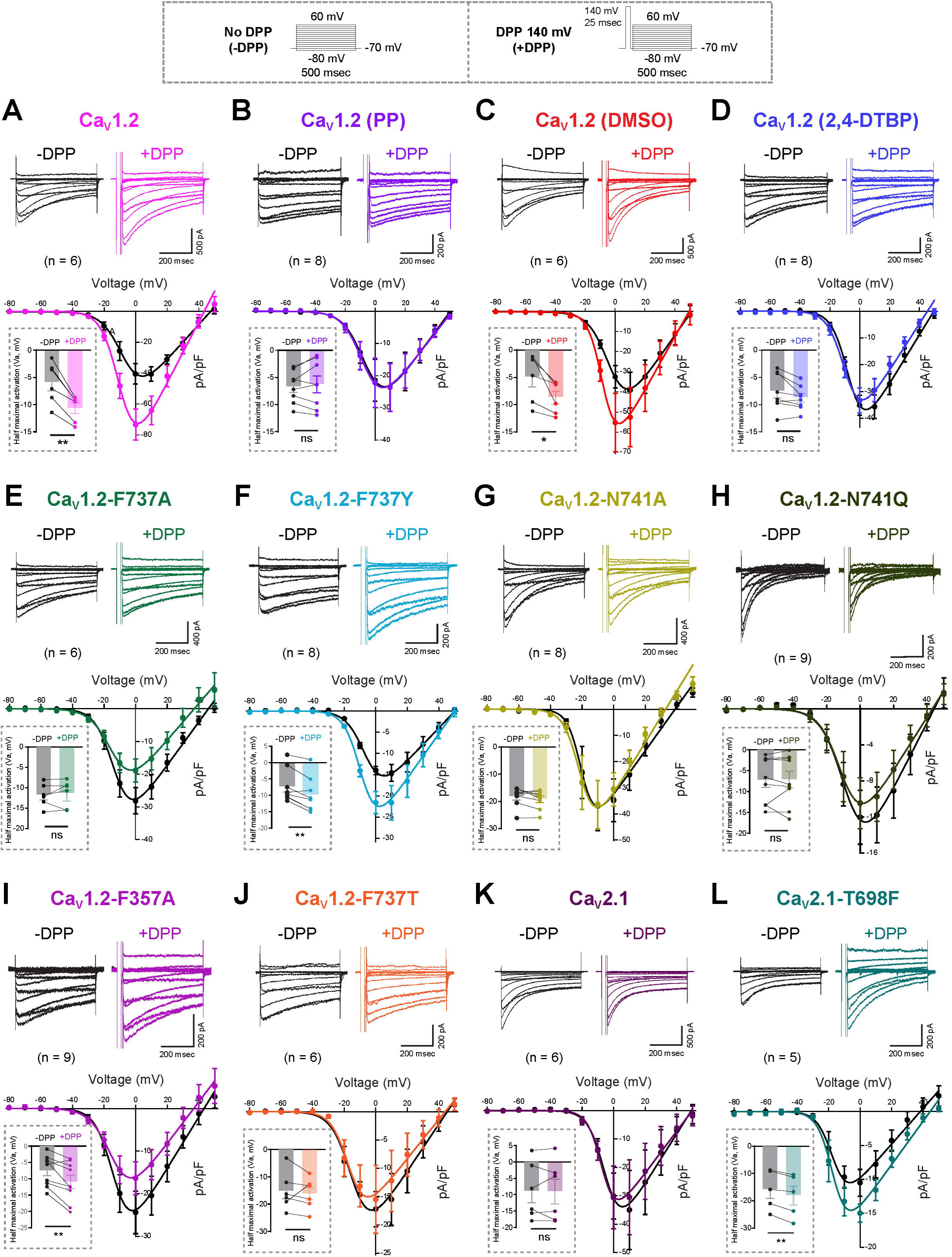
DPP-induced shift in voltage-dependent activation of L-type channels: **(A-L)** Representative whole-cell Ba² current traces of WT and mutant Ca_V_1.2 and Ca_V_2.1 channels in various conditions obtained from -80 to +60 mV voltage steps of 500 msec, with (+DPP, colored) or without (−DPP, black) a DPP of +140 mV for 25 msec, are shown in the upper panels. The voltage protocol used is shown in the dotted box above. Corresponding current-voltage (I-V) plots are shown below each respective current trace, with current density (pA/pF) plotted as a function of different test-potential pulses. Inset bar plots show the half-maximal activation (V_a_) voltages for the specified channels. All bar plots represent mean ± SEM. Experimental replicate cell numbers are indicated in brackets. An unpaired Student’s t-test is used to determine statistical significance. *p ≤ 0.05, **p ≤ 0.01, ***p ≤ 0.001, ****p < 0.0001, ns > 0.05; ns, non-significant.

In analogy to whole-cell current recordings, a transition from low to high open-channel probability was observed in single-channel recordings of Ca_V_1.2 in response to DPP stimulus^13,15,17,25,26^. A strong DPP of +100 mV or greater is reported to shift the equilibrium between the opening and closing rate constants toward longer-duration openings, thereby increasing open probability and mean open times. To validate this, cell-attached single-channel recordings of Ca_V_1.2 channels were performed in tsA-201 cells coexpressing α_1C_ and β_1b_ subunits, which exhibited the highest VDF in whole-cell recordings. At a 0 mV holding potential, we recorded brief opening and closing events of single Ca_V_1.2 channels (***Fig. 1E, black traces***), with a mean amplitude of ∼1.6 pA in glass-bottle-stored external solutions, in agreement with earlier observations^44^. A brief 100-msec DPP to +100 mV elicited comparatively longer openings (***Fig. 1E, red traces***) that are reflected by a significant increase in the mean open probability (NP_o_) (0.07 ± 0.03 to 0.10 ± 0.03, p = 0.0156) (***Fig. 1F***). Open dwell time analysis with a nonlinear least-squares fitting of single channel traces yielded three exponentials with open time constants (τ_o_) of ∼1.3, 7.8, and 54.6 msec representing short, medium, and long open states (***Fig. 1G***). DDP stimulation to +100 mV significantly increased channel occupancy, specifically in the long-open state (2.4 ± 0.8% to 8.79 ± 1.9%, p = 0.007) without affecting the short or medium open states (***Fig. 1G***), implying a shift in gating modes^13,17,25^. In contrast, no change was observed in NP_o_ and open dwell time states of Ca_V_1.2 in cells bathed in external solutions stored in PP tubes for 5-7 days in response to a 100 mV DPP stimulus (***Fig. 1H-1J***), which strongly correlates with the whole-cell observations of VDF inhibition. Additionally, no measurable difference was also observed in the basal occupancy of channels in the long-open state among cells bathed in glass-stored solutions and PP-stored solutions (***Fig. 1G and IJ***). These results indeed suggest VDF inhibition by PP-stored solutions rather than channels entering into a pre-facilitated state in response to DPP stimulus.

Altogether, whole-cell and single-channel investigations suggested that PP-stored solutions inhibit L-type channel VDF, implying a probable role of plastic leachables, without affecting normal gating processes.

### Characterization of leachables in PP stored solutions

Stabilizers, antioxidants, and plasticizers, which are routinely added during the manufacturing of PP materials, continuously leach into aqueous solutions over time, posing environmental hazards and causing diverse pathophysiological effects, ranging from endocrine disruption to developmental and cognitive dysfunctions^40–42,45,46^. Our initial observations with external solutions stored in PP tubes, which completely abolished the L-type channel VDF, guided us to characterize the leachables from PP-incubated buffer solutions and investigate their effects on the VDF phenomenon of Ca_V_1.2 and Ca_V_1.3 channels.

DD water or a 1M aqueous BaCl stock solution was incubated in laboratory PP tubes for 1-2 weeks. Subsequently, the leached compounds were extracted by liquid-liquid extraction with HPLC grade ethyl acetate and analyzed by gas chromatography-mass spectrometry (GC-MS) on a triple quadrupole GC-MS (***Fig. 3***). The retention time (RT) and mass-to-charge (*m/z*) ratios of the GC-MS peaks indicated the presence of two tentative compounds 1,3-di-tert-butylbenzene (1,3-DTBB) and 2,4-DTBP (***Fig. 3B***), as referenced in the National Institute of Standards and Technology (NIST) mass spectral database^46–48^, and performed further characterization. The presence of both 1,3-DTBB and 2,4-DTBP peaks was observed in extracts from DD water and BaCl -stored PP vials, with extracted ion chromatograms (EICs) at 174.6 and 190.7 *m/z*, respectively, and RTs of 14.2 and 17.6 min, respectively (***Fig. 3B***). The leaching of 1,3-DTBB and 2,4-DTBP from PP tubes was validated by comparing the RT and mass spectra (MS) fragment *m/z* ratio at EIC 174.6 and 190.7 m/z, respectively, with the commercial standards of these compounds under GC-MS examination (***Fig. 3B-3F***). In similar experimental conditions, no peaks of 2,4-DTBP or 1,3-DTBB were observed in extracts of DD water or aqueous BaCl_2_ incubated in glass reagent bottles (***Fig. 3B***). Additionally, thin-layer chromatography (TLC) enrichment followed by liquid chromatography-mass spectrometry (LC-MS) analysis identified another leachable compound, di-2-ethylhexyl phthalate (DEHP) in extracted smaples of aqueous 1M BaCl_2_ stored in PP tubes for a longer duration of 1 month, with an EIC at 391.2 and an RT of 16.1 min (***Fig. S6A-S6C)***. This was further confirmed by analyzing the RT and MS-MS fragment *m/z* ratios relative to commercial standards (***Fig. S6C and S6D***).

**Figure 3:**
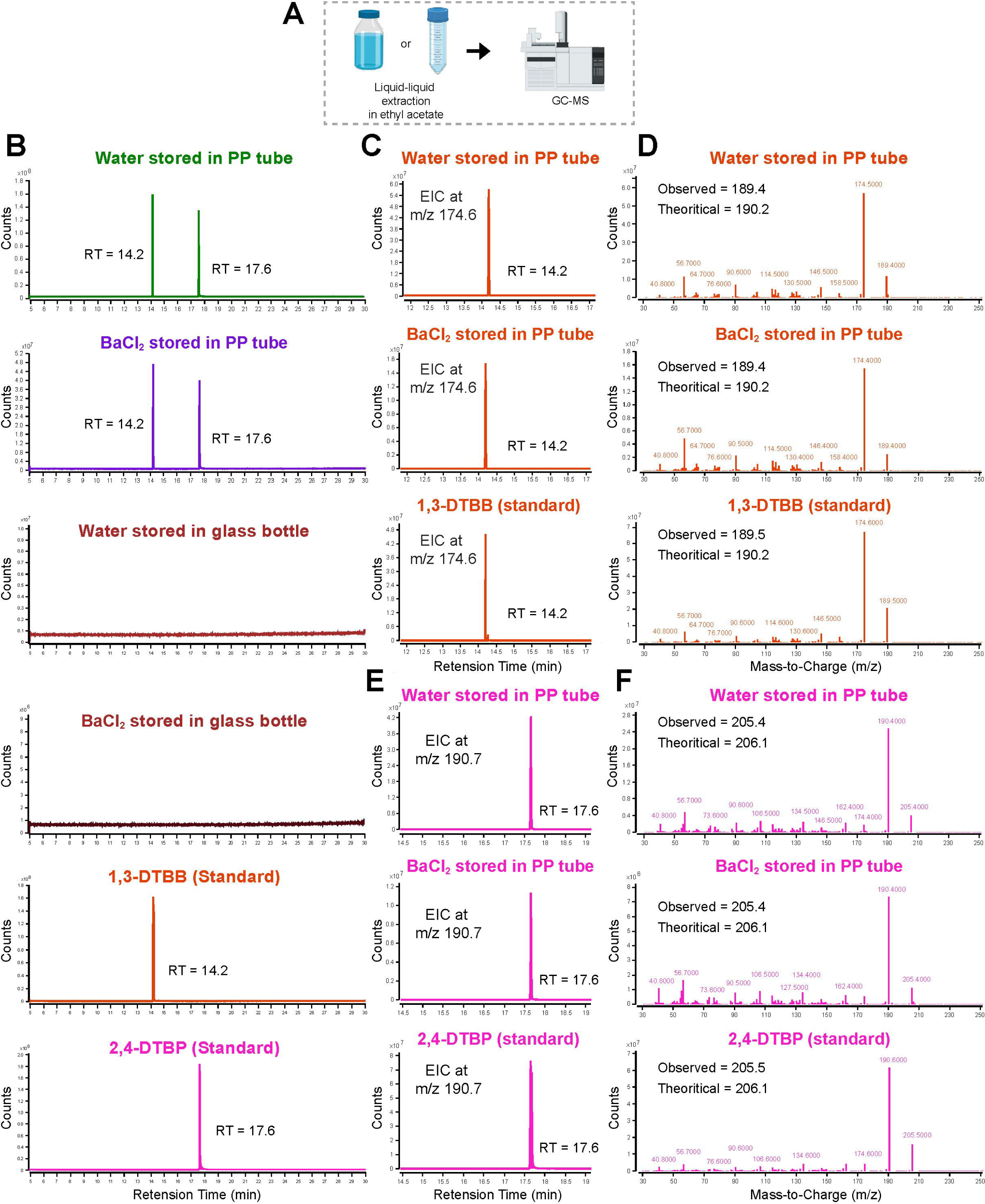
GC-MS analysis confirms leaching of 1,3-DTBB and 2,4-DTBP in PP-stored solutions: **(A)** Schematic representation of GC-MS analysis of liquid-liquid extracted water samples stored either in glass bottles or PP tubes. **(B)** GC-MS comparative chromatogram extract of DD water or 1 M aqueous BaCl_2_ stored in PP tubes versus glass bottles in HPLC-grade ethyl acetate. Two distinctive peaks appeared in the chromatograms of DD water and BaCl_2_ extracts stored in PP tubes. No peak was observed in DD water and BaCl_2_ extracts stored in glass bottles. GC-MS chromatograms of the commercial analytical standards for 1,3-DTBB and 2,4-DTBP are shown below. **(C)** EIC at 170.6 m/z of DD water, aqueous 1M BaCl_2_ extract stored in PP tubes, and the analytical standard of 1,3-DTBB in ethyl acetate displaying a retention time of 14.2 min. **(D)** MS/MS spectra or Mass-spectra at EIC 170.6 m/z of DD water, aqueous 1M BaCl_2_ stored in PP tubes, or the analytical standard 1,3-DTBB, indicating the theoretical and observed mass in KD. **(E)** EIC at 190.7 m/z of DD water, aqueous 1M BaCl_2_ extract stored in PP tubes, and the analytical standard of 2,4-DTBP in ethyl acetate displaying a retention time of 17.6 min. **(F)** MS/MS spectra or Mass-spectra at EIC 190.7 m/z of DD water, aqueous 1M BaCl_2_ stored in PP tubes, or the analytical standard 2,4-DTBP, indicating the theoretical and observed mass in KD.

Together, GC-MS and LC-MS facilitated the sensitive detection and characterization of three common plastic additive compounds—2,4-DTBP, 1,3-DTBB, and DEHP in BaCl_2_ and water-incubated samples in PP tubes.

### 2,4-DTBP potently inhibits L-type channel VDF

To test the possible inhibitory effect of the identified compounds on L-type channel VDF, whole-cell patch clamp recordings were performed in cells coexpressing either Ca_V_1.2 or Ca_V_1.3 cDNAs along with β_1b_ and exposed to 1 µM of either of the compounds, dissolved in DMSO solvent. At +120 mV DPP, Ca_V_1.2 and Ca_V_1.3 channels coexpressed with β_1b_ exhibited a VDF of 64.86 ± 9.19%, n = 9, and 35.46 ± 6.12%, n = 8, respectively, in the presence of DMSO control (0.01%) (***Fig. 4A, 4B, and S7***). However, VDF of Ca_V_1.2 and Ca_V_1.3 channels remained unaffected by 1 µM of either 1,3-DTBB or DEHP (***Fig. S7A-S7D***) (***Table S1***). Notably, 1 µM 2,4-DTBP completely inhibited Ca_V_1.2 and Ca_V_1.3 VDF (5.17 ± 4.15%, n = 8, p < 0.001; 4.54 ± 1.35%, n = 7, p < 0.001 at 120 mV DPP, repectively) (***Fig. S7A–S7D***). Furthermore, VDF measurements in cells coexpressing β_1b_ with Ca_V_1.2 or Ca_V_1.3 cDNAs revealed concentration-dependent 2,4-DTBP blockade (100, 250, or 500 nM), achieving complete inhibition at 500 nM (2–3% VDF at 120 mV; ***Fig. 4A and 4B***). A similar inhibition was observed in cells coexpressing α_2_δ_1_and β_1b_ (***Fig. S9A and S9B***). Further, no change in Ca_V_1.2 channel V_a_ was observed upon stimulation with DPP at +140 mV in the presence of 500 nM 2,4-DTBP, which was evident in control DMSO solutions (***Fig. 2C and 2D***). Also, in paired recordings, exchanging control solutions with solutions containing 500 nM 2,4-DTBP for 5-10 min completely blocked the VDF of Ca_V_1.2 in response to +140 mV DPP without affecting the basal current density (***Fig. S5C and S5D***), implying inhibition of VDF. Despite complete VDF inhibition at 500 nM 2,4-DTBP, no significant change was observed in the current density, voltage dependence, inactivation kinetics, and residual R_400_ current pattern of Ca_V_1.2 and Ca_V_1.3 channels coexpressed with β_1b_ (***Fig. S8)*** or with β_1b_and α_2_δ_1_(***Fig. S9C and S9D) (Table S1***), thereby complementing PP-incubated solutions’ VDF outcomes.

**Figure 4:**
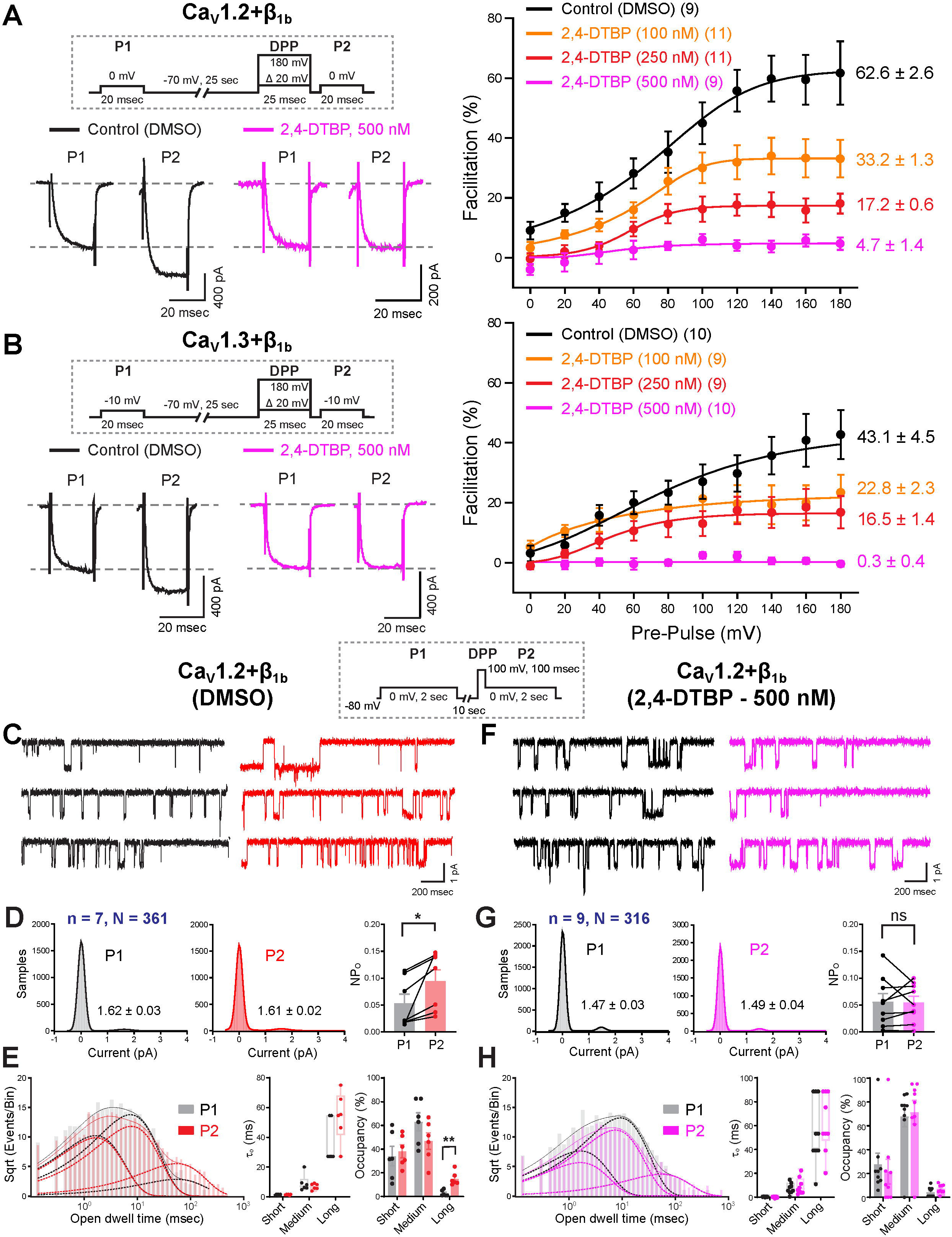
2,4-DTBP potently inhibits the VDF of L-type channels: **(A and B)** Representative whole-cell current traces of L-type Ca_V_1.2 **(A)** and Ca_V_1.3 **(B)** channels coexpressed with β_1b_ subunit in tsA-201 cells recorded in external solutions containing 500 nM 2,4-DTBP (magenta) or DMSO (black) before (P1) or after (P2) the DPP to 100 mV. Mean fitted plots (right) show maximum facilitation of Ca_V_1.2 **(A)** and Ca_V_1.3 **(B)** channel currents with 100, 250, or 500 nM 2,4-DTBP or DMSO control in response to DPP ranging from 0 to 180 mV. The voltage protocol is shown in the dotted box above. Experimental replicate cell numbers are indicated in brackets. **(C and F)** Exemplary cell-attached single-channel current traces of Ca_V_1.2 coexpressed with β_1b_subunit in tsA-201 cells recorded in the presence of DMSO **(C)** or 500 nM 2,4-DTBP **(F)**. The voltage protocol used is shown in the dotted box above. **(D and G)** The histogram plots show the single-channel current amplitudes, while the bar plots display the open probability of Ca_V_1.2 cell-attached recordings obtained at 0 mV step of 2 sec in the presence of DMSO **(D)** or 500 nM 2,4-DTBP **(G)** before (P1) or after (P2) the DPP to 100 mV for 100 msec. N = number of cells, n = number of single-channel traces. A non-parametric paired Student’s t-test is used to determine statistical significance. **(E and H)** The logarithmic histogram plots show the open dwell-time distributions of Ca_V_1.2 channels in DMSO or 2,4-DTBP-containing solutions, fitted by nonlinear least squares, while the bar plots display open time constants and their occupancy in short, medium, and long open states. One-way ANOVA with pairwise comparison was used to determine statistical significance. All plots represent mean ± SEM. *p ≤ 0.05, **p ≤ 0.01, ***p ≤ 0.001, ****p < 0.0001, ns > 0.05; ns, non-significant.

Further examination of the mode of L-type channel VDF inhibition was performed by recording Ca_V_1.2 single-channel currents in the presence of 500 nM 2,4-DTBP or an equal volume of DMSO used as a control. Ca_V_1.2 channels coexpressed with β_1b_ displayed a significant increase in the NP_o_ (0.05 ± 0.02 to 0.09 ± 0.02, p = 0.0156) and an increment in open dwell time, specifically the long open state (2.74 ± 1.01% to 14.98 ± 2.3%, p = 0.0098) upon stimulation with a DPP of +100 mV in DMSO-exposed cells (***Fig. 4C-4E***), implying a shift in gating due to VDF. In contrast, no significant change in NP_o_ or open dwell time states was observed in cells incubated with 500 nM 2,4-DTBP in response to a DPP of +100 mV (***Fig. 4F-4H***), indicating inhibition of the mean open time gating shift, consistent with single-channel observations in PP-incubated solutions.

Collectively, whole-cell and single-channel analyses illustrate the specific inhibition of the VDF of Ca_V_1.2 and Ca_V_1.3 L-type channels by the common plastic stabilizer 2,4-DTBP.

### L-type channel VDF inhibition in hippocampal neurons

L-type channel VDF is known to regulate the frequency of action potentials in response to stronger depolarizing stimuli in neurons and cardiomyocytes^16,29,49^. To test the results obtained in expression systems, VDF measurements were carried out in cultured hippocampal pyramidal neurons at DIV 7-9, during which L-type Ca_V_1.2 and Ca_V_1.3 channel isoforms are abundantly expressed in the soma, proximal dendrites, and postsynaptic regions^15,17,49^. Whole-cell voltage clamp steps to 0 mV of 20 msec from a holding potential of -70 mV produced an inward mean current of 255.63 ± 38.59 pA (n = 11), which was facilitated to 319.32 ± 43.57 pA following the DPP to 100 mV for 25 msec, accounting for 27.84 ± 3.26% VDF, in 5 mM BaCl_2_ containing ACSF stored in glass bottles (***Fig. S10***). However, the magnitude of VDF was drastically reduced to 3.33 ± 2.56% (n = 6), p = 0.0001, in neurons bathed in ACSF prepared with PP stored DD water, and to 3.84 ± 1.75% (n = 10), p <0.0001, in ACSF containing 500 nM 2,4-DTBP (***Fig. S10***). These results demonstrate that PP-stored solutions and 2,4-DTBP block L-type channel VDF in both exogenous and endogenous systems.

### 2,4-DTBP docks at the DI–DII PD interface

Specific blockade of L-type channel VDF by the plastic stabilizer 2,4-DTBP suggests a possible interaction and steric hindrance that restricts the conformational rearrangements crucial for DPP-induced VDF in L-type channels. In an effort to identify the molecular determinants of VDF, we probed the binding site of 2,4-DTBP on L-type channels by performing docking followed by MD simulations using the reported Cryo-EM structures of human Ca_V_1.2 (PDB:8WE6)^50^ and Ca_V_1.3 (PDB:7UHG)^51^ protein complexes as templates. Docking with Autodock 4.2 predicted a strong interaction of 2,4-DTBP with the α_1_ subunit of Ca_V_1.2 (***Fig. 5A and 5B***) and Ca_V_1.3 (***S10A and S10B***) in the DI–DII PD fenestration pocket, with binding energies of -5.25 kcal/mol and -4.66 kcal/mol, respectively. All-atom MD simulations for 500 ns exhibited stable positioning of 2,4-DTBP at the DI–DII PD interface of Ca_V_1.2 (***Fig. 5F and Video S1)*** and Ca_V_1.3 (***S11F and Video S2)***, with RMSD of ∼2-4 Å in five different seeds, ensuring varied random initial velocities. The protein backbone of Ca_V_1.2 (***Fig. 5D***) and Ca_V_1.3 (***Fig. S11D)*** stabilized with an RMSD of <4 Å. The phenol ring, along with the butyl side chains of 2,4-DTBP, was stabilized in a hydrophobic pocket formed by residues of DIS6, DIS5, DIP1, and DIIS6 helices in Ca_V_1.2 (***Fig. 5B***) and Ca_V_1.3 (***S11B),*** constituting the binding interface. The distance between the center of mass (CoM) of 2,4-DTBP and the binding interface of Ca_V_1.2 (***Fig. 5G***) and Ca_V_1.3 (***Fig. S11G)*** exhibited an average RMSD of ∼2 Å, demonstrating stable binding. Moreover, the 2,4-DTBP binding pocket was reported to be in an open conformation that accommodates a blocking lipid in the Ca_V_1.2^5^ and Ca_V_1.1^6,52^ Cryo-EM structures.

**Figure 5:**
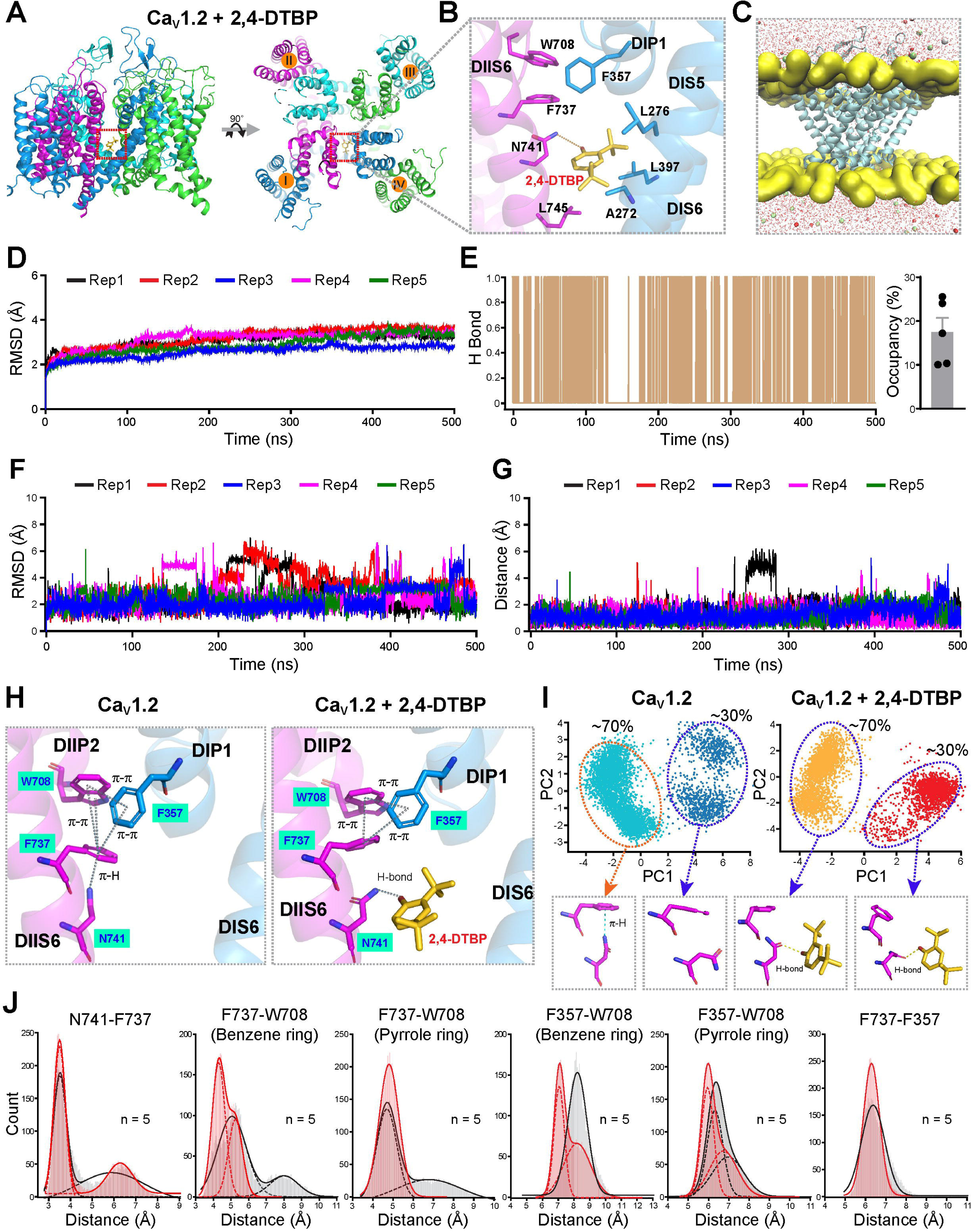
2,4-DTBP disrupts the π-H and π-π quadrangle interaction at the DI and DII PD interface of the Ca_V_1.2 (α_1C_) subunit. **(A)** Side view (left) and top view (right) of the human Ca_V_1.2 protein backbone displaying 2,4-DTBP binding at the DI and DII PD interface. **(B)** Zoomed view of the 2,4-DTBP binding pocket at the DI-DII fenestrations. Key residues forming hydrophobic and H-bonding with 2,4-DTBP in the DIS6, DIS5, and DIIS6 are indicated. **(C)** Graphical representation of the MD simulation setup of the Ca_V_1.2 pore domain with or without 2,4-DTBP. **(D)** Root-mean-square deviation (RMSD) of Ca_V_1.2 backbone during five independent 500 ns MD simulations, showing overall structural stability of the protein-ligand complex. **(E)** Exemplary hydrogen bond occupancy between N741 and 2,4-DTBP during the 500 ns MD-simulation. The bar plot shows the average hydrogen-bond occupancy across five replicates. **(F)** RMSD of 2,4-DTBP in the binding interface during 500 ns MD simulations across five independent replicates, demonstrating ligand stability. **(G)** Distance between the center of mass of 2,4-DTBP and the binding pocket of Ca_V_1.2 in the DI-DII PD interface over the course of 500 ns MD simulations across five independent replicates. **(H)** A zoomed view of the DI and DII PD interface of Ca_V_1.2 reveals feasible π-H and π-π quadrangle interactions among the F357, F737, W708, and N741 residues. **(I)** The PCA of the side chains of F737 and N741 reveals distinct clusters of possible conformations. The representative conformational orientations for each cluster are shown below. **(J)** Centroid-to-centroid distances between the selected residues of Ca_V_1.2 in the absence and presence of 2,4-DTBP over the simulation trajectories.

### 2,4-DTBP interferes with an asymmetric π-H and π-π quadrangle

Based on electrophysiological and computational evidence, it is likely that 2,4-DTBP sterically interferes with the conformational rearrangements that operate for the DPP-induced VDF. To characterize this, comparative MD simulations of both Ca_V_1.2 and Ca_V_1.3 α_1_ subunits were performed in the absence (apo) and presence (holo) of 2,4-DTBP in five different replicates for 500 ns. Interestingly, we observed a favorable geometry for feasible π-H bonding between the benzene ring of F737 and the amide side chain of N741 in DIIS6 in Ca_V_1.2 apo simulations (***Fig. 5H***). Also, feasible π-π interactions were observed between the benzene rings of F737, W708, and F357 located at the DIIS6, DIIP2, and DIP1 helices, respectively. These interactions together constitute an interdomain quadrangle at the DI– DII PD interface (***Fig. 5H***). An equivalent quadrangle composed of a feasible π-H interaction between F736 and N740 in DIIS6, and π-π interactions between F736, W707, and F358 AAs located at similar DIIS6, DIIP2, and DIP1 helices were observed in Ca_V_1.3 (***Fig. S11H***). A principal component analysis (PCA) of the sidechains, followed by K-means clustering across five independent replicates in apo simulations, revealed that ∼70% of sidechains have favorable orientations for establishing a feasible π-H bond between F737 and N741 in Ca_V_1.2 (***Fig. 5I***) and between F736 and N740 in Ca_V_1.3 (***Fig. S11I***). However, favorable orientation clusters for π-H bonding were not observed in both Ca_V_1.2 and Ca_V_1.3 holo simulations in the presence of 2,4-DTBP. This decrease is due to the constant engagement of the 2,4-DTBP’s hydroxyl group in a transient hydrogen bond with the sidechain of N741 in Ca_V_1.2 (***Fig. 5E and Video S1***) and N740 in Ca_V_1.3 (***Fig. S11E and Video S2***), with an average occupancy of ∼15-18%. Notably, this reorientation of asparagine in both Ca_V_1.2 and Ca_V_1.3 leads to a closer association of W708 with F737 and F357 in Ca_V_1.2 (***Fig. 5J***)) and between W707 with F736 and F358 in Ca_V_1.3 (***Fig. S11J***) as reflected in the centroid-centroid distance analysis in holo simulations compared to apo. This effectively destabilized the overall π-H and π-π quadrangle geometry.

Altogether, the docking and MD simulation results suggested a favorable confirmation feasible for forming the π-H and π-π interactions in the DI-DII PD fenestration of Ca_V_1.2 and Ca_V_1.3 L-type channels, which are sterically hindered in the presence of 2,4-DTBP.

### Asymmetric π-H and π-π quadrangle contribute to VDF

The disruption of π-H and π-π quadrangle association by 2,4-DTBP indicated a crucial contribution of the quadrangle-forming residues in DPP-induced VDF of L-type channels. To determine the molecular endpoints of VDF, a series of point mutations was generated in the Ca_V_1.2 α_1_ subunit, including F737A, N741A, F357A, and W708A, which could destabilize the π-H and π-π quadrangle interactions (***Fig. 6A***)). Whole-cell VDF measurements were performed by coexpressing the point mutants along with β_1b_. A Complete disappearance of VDF was recorded in Ca_V_1.2-F737A and Ca_V_1.2-N741A mutants across DPPs ranging from 0-180 mV, while Ca_V_1.2-F357A and Ca_V_1.2-W708A channels showed intermediate 15-20% VDF (***Fig. 6B-6D***). Further, no shift in V_a_ was observed in Ca_V_1.2-F737A and Ca_V_1.2-N741A channels (***Fig. 2E and 2G***), in response to a DPP of +140 mV. While Ca_V_1.2-F357A channels displaying intermediate facilitation showed a significantly hyperpolarizing shift in V_a_ (***Fig. 2I***)). Notably, the elimination of VDF and hyperpolarizing shift in V_a_ could be completely rescued by introducing a tyrosine in place of F737, which effectively restores the π-H and π-π quadrangles (***Fig. 6A and 2F***). The resultant Ca_V_1.2-F737Y channels exhibited 54.83 ± 10.12% (n = 7) VDF in response to 120 mV DPP, which was comparable to the control Ca_V_1.2 channels displaying ∼61% VDF (***Fig. 6B-6D***). In contrast, replacing asparagine with glutamine at position 741 did not support the π-H and π-π quadrangle interactions due to its longer side chain, and the resultant channel Ca_V_1.2-N741Q completely lacked VDF (***Fig. 6A-6D***) and a hyperpolarizing shift in V_a_, in response to DPP (***Fig. 2H***)). The whole-cell observations were further verified by single-channel analysis, which demonstrated a complete loss of DPP-induced VDF in the Ca_V_1.2-F737A mutant (***Fig. 6E-6G***). While Ca_V_1.2-F737Y channels exhibited a significant increase in NP_o_ (0.13 ± 0.04 to 0.19 ± 0.04, p = 0.0156) and a shift in open dwell time, specifically the long state (10.15 ± 3.72 to 15.60 ± 4.7, p = 0.015) in response to 100 mV DPP compared to the openings recorded without prestimulus (***Fig. 6H-J***)). However, a comparatively higher open dwell time was observed in Ca_V_1.2-F737Y channels compared to Ca_V_1.2-WT (***Fig. 1E and 1G***, p <0.0001). Additionally, Ca_V_1.2-F737A channels, which lack VDF, showed significantly greater inactivation than Ca_V_1.2-WT^53^, a phenotype rescued in Ca_V_1.2-F737Y channels (***Fig. S11C, Table S1***). Together, these results demonstrate the crucial role of π-H and π-π quadrangle interactions at the DI–DII PD interface in eliciting VDF in addition to channel activation and inactivation gating.

**Figure 6:**
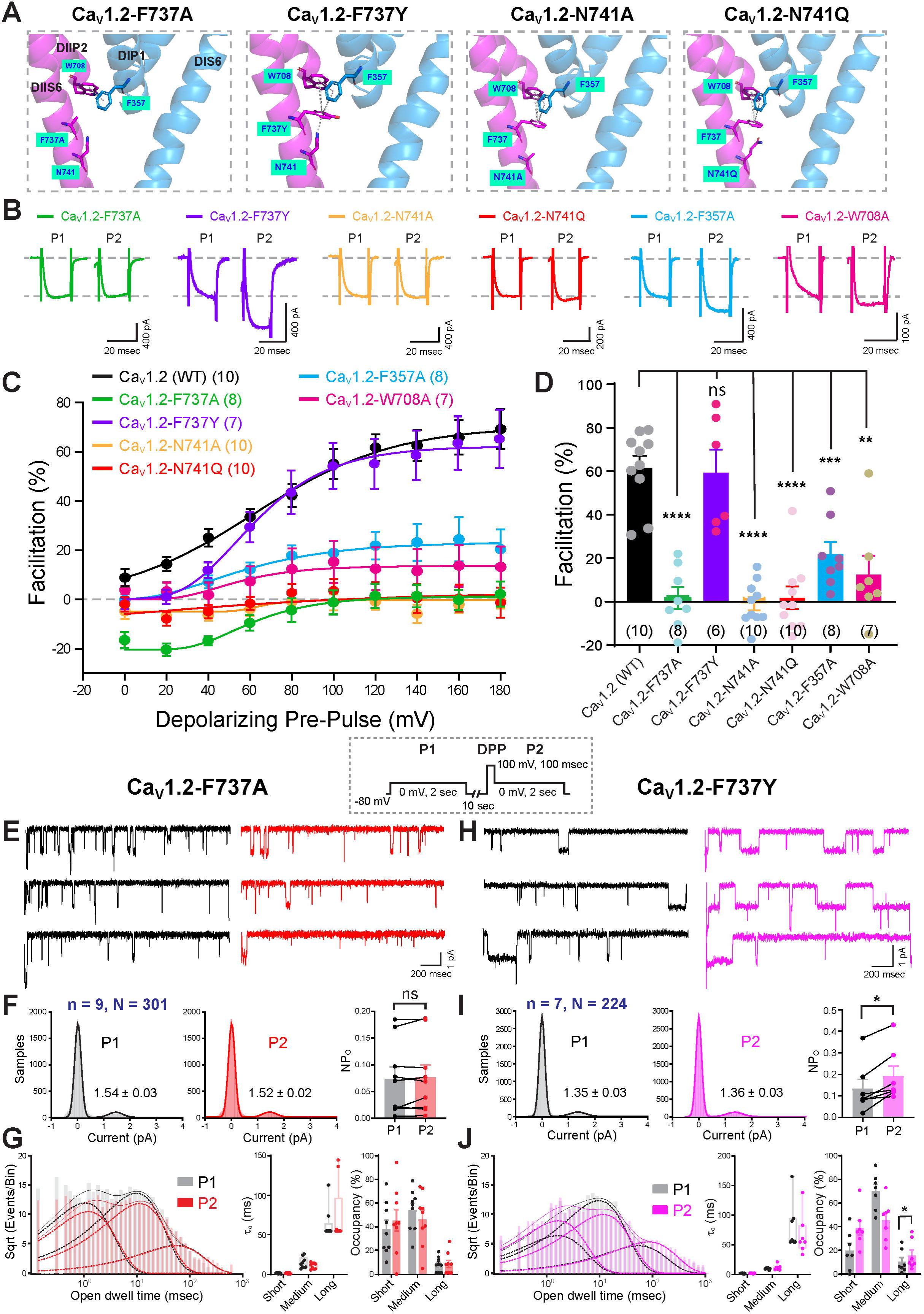
π-H and π-π quadrangle interactions within DI-DII PD fenestration contribute to Cav1.2 channel VDF. **(A)** Schematic of DI–DII PD fenestration in Ca_V_1.2 WT, F737A, F737Y, N741A, and N741Q mutants generated from homology modeling. Possible π-H and π-π quadrangle interactions are shown as dotted lines. **(B)** Representative whole-cell current traces of Ca_V_1.2 point mutants at DI–DII PD fenestration coexpressed with β_1b_ in tsA-201 cells before (P1) or after (P2) DPP to 100 mV. **(C)** Mean fitted plots of Ca_V_1.2 WT and DI–DII PD mutants coexpressed with β_1b_in response to DPP ranging from 0 to 180 mV. **(D)** Bar plots depict maximum VDF at 120 mV DPP for Ca_V_1.2 WT and DI–DII PD mutants. One-way ANOVA was used to determine statistical significance comparing the mutants with the WT. **(E and H)** Exemplary cell-attached single-channel current traces of Ca_V_1.2-F737A **(E)** and Ca_V_1.2-F737Y **(H)** channels coexpressed with β_1b_ in tsA-201 cells before and after the DPP to 100 mV. The voltage protocol used is shown in the dotted box. **(F and I)** The histogram plots show the single-channel current amplitudes, while the bar plots display the open probability of Ca_V_1.2-F737A **(F)** and Ca_V_1.2-F737Y **(I)** at 0 mV step of 2 sec before (P1) or after (P2) the DPP to 100 mV for 100 msec. N = number of cells, n = number of single-channel traces. A non-parametric paired Student’s t-test is used to determine statistical significance. **(G and J)** The logarithmic histogram plots show the open dwell-time distributions of Ca_V_1.2-F737A **(G)** and Ca_V_1.2-F737Y **(J)** channels fitted by nonlinear least squares, while the bar plots display open time constants and their occupancy in short, medium, and long open states. One-way ANOVA with pairwise comparison was used to determine statistical significance. All plots represent mean ± SEM. Experimental replicate cell numbers are indicated in brackets. *p ≤ 0.05, **p ≤ 0.01, ***p ≤ 0.001, ****p < 0.0001, ns > 0.05; ns, non-significant.

In contrast to the DI–DII PD interface, similar π-H and π-π quadrangle interactions were not observed at other PD fenestrations, suggesting asymmetric positioning. Mutations in AA located at similar positions to F737, F357, and W708 in the Ca_V_1.2 DII–DIII, DIII–DIV, and DI–IV PD interface to alanine resulted in an intermediate VDF in response to DPPs (***Fig. S13***). But a complete inhibition was not observed in any of the mutants.

MD simulations and mutagenesis studies collectively show that asymmetric π-H and π-π quadrangle interactions at the DI–DII PD interface serve as key molecular endpoints that drive VDF in L-type channels.

### Engineering VDF on Ca_V_2.1 channels

DPP-induced VDF is a unique feature of L-type Ca_V_1 channels, while Ca_V_2.1-2.3 HVA channels exhibit G_βγ_ tonic inhibition relief during repeated depolarizations, distinct from L-type VDF^54,55^. Further, Ca_V_2.1 channels didn’t elicit facilitation in P2 currents compared with the P1 voltage step, in response to a DPP voltage step of 0-180 mV (***Fig. 7C-7E***). To understand this deviation, Multiple sequence alignment (MSA) was performed between the helices constituting DI and DII PD interface of Ca_V_1.1-1.4 channels with that of Ca_V_2.1-2.3 channels, with special emphasis on the AAs participating in the π-H and π-π quadrangle interaction. Ca_V_2.1-2.3 channels share similar residues at three positions but have threonine instead of the critical central phenylalanine required for the π-H and π-π quadrangle geometry (***Fig. 7A and S14A***). To further test the central role of π-H and π-π quadrangle interactions at the DI–DII PD interface in VDF, we generated a Ca_V_2.1-T698F mutant that resembles the Ca_V_1 channel architecture. Indeed, the Ca_V_2.1-T698F channel mutation proved to be sufficient to induce a strong VDF in response to DPP of 0-180 mV, which was absent in Ca_V_2.1-WT channels (54.84 ± 8.12%, n = 8, from 3.35 ± 1.50%, n = 7, respectively, at 180 mV DPP, p < 0.0001) (***Fig. 7C-7E***). By comparison, substitution of central phenylalanine with threonine in Ca_V_1.2-F737T channels resulted in the complete abolition of the VDF, as assessed by whole-cell and single-channel recordings (***Fig. 7C-7E and S15***). Importantly, Ca_V_2.1-T698F channels displaying VDF also exhibited a ∼-3 mV significant hyperpolarizing shift of V_a_ in response to 140 mV DPP, which was absent in Ca_V_2.1 WT channels (***Fig. 2K and 2L***). Alternatively, Ca_V_1.2-F737T lacking VDF didn’t show any significant shift in V_a_ due to DPP (***Fig. 2J***)).

**Figure 7:**
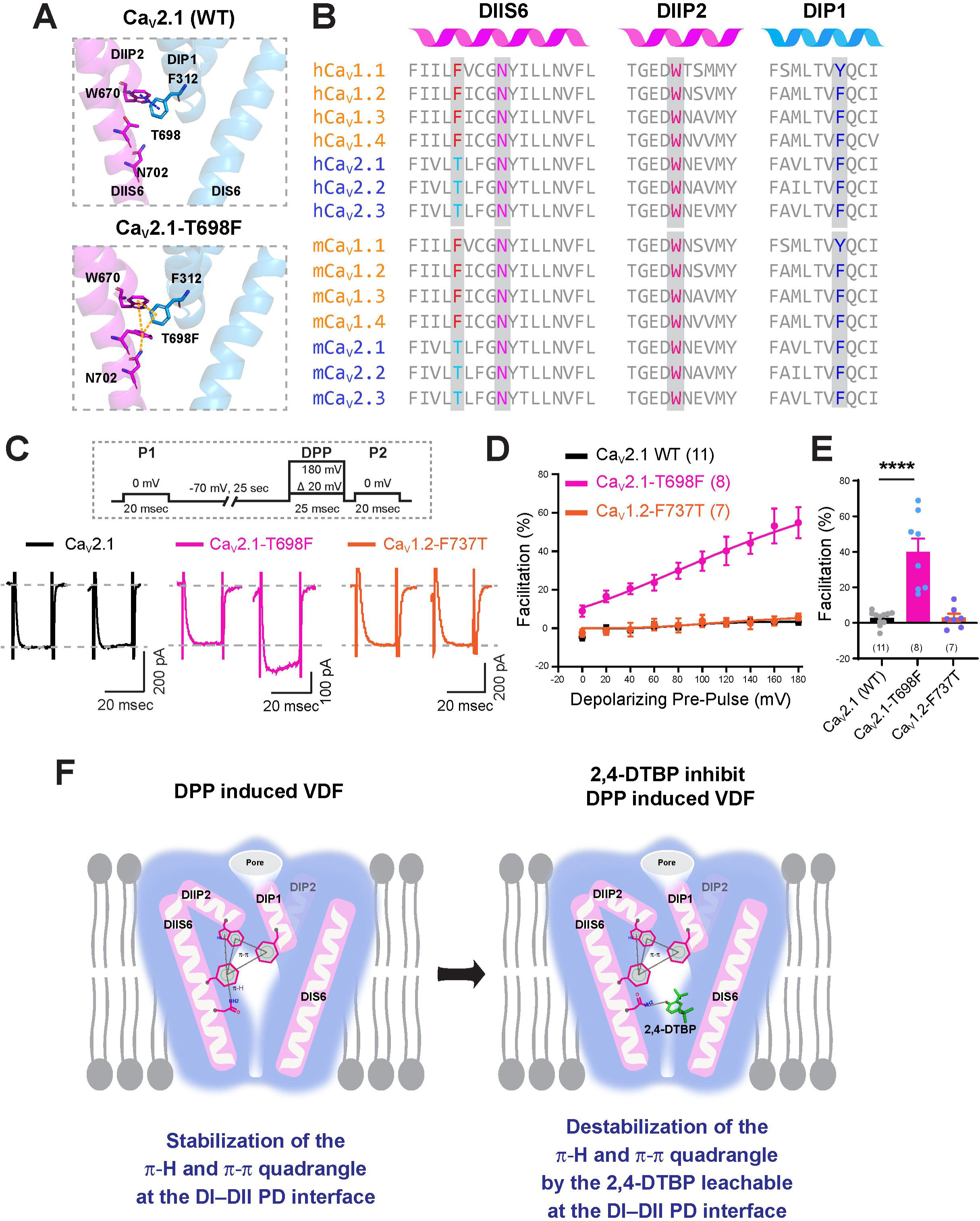
Incorporation of π-H and π-π quadrangle interactions at the DI-DII PD interface elicits VDF in Ca_V_2.1 channels. **(A)** Structural representation of DI–DII PD interface in Ca_V_2.1-WT and Ca_V_2.1-T698F mutant. Possible π-H and π-π quadrangle interaction in Ca_V_2.1-T698F mutant shown as dotted lines. **(B)** Aligned amino acid sequences of DIIS6, DIIP2, and DIP1 PD helices from human (h) and mouse (m) L-type (Ca_V_1.1–1.4) and Ca_V_2 (2.1–2.3) channels. Conserved and unique residues participating in π-H and π-π quadrangle interactions are highlighted. **(C)** Representative whole-cell current traces of Ca_V_2.1 (WT), Ca_V_2.1-T698F, and Ca_V_1.2-737T point mutant channels coexpressed with β_1b_ subunit in tsA-201 cells before (P1) or after (P2) the DPP to 100 mV. The voltage protocol is represented in the dotted box above. **(D)** Mean fitted plots of Ca_V_2.1-WT, Ca_V_2.1-T698F, and Ca_V_1.2-F737T mutants in response to DPP ranging from 0 to 180 mV. **(E)** Bar plots display the maximum VDF at 120 mV DPP for Ca_V_2.1-WT, Ca_V_2.1-T698F, and Ca_V_1.2-F737T mutants. Unpaired Student’s t-test used statistical significance. **(F)** Graphical representation of the π-H and π-π quadrangle interactions at the DI and DII PD interface in L-type Ca_V_ channels. DPP voltages stabilize the π-H and π-π quadrangle interaction. Conversely, 2,4-DTBP leachable destabilizes the crucial π-H and π-π quadrangle interactions by forming an H-bond with the participating asparagine and thereby inhibiting the VDF. All plots represent mean ± SEM. Experimental replicate cell numbers are indicated in brackets. *p ≤ 0.05, **p ≤ 0.01, ***p ≤ 0.001, ****p < 0.0001, ns > 0.05; ns, non-significant.

These results collectively highlight the crucial role of π-H and π-π quadrangle interactions, asymmetrically positioned at the DI–DII PD interface of the Ca_V_1 L-type channels, which constitute the molecular endpoint of VDF. These interactions can also be transferred to non-VDF Ca_V_2 channels, enabling robust DPP-induced facilitation.

## Discussion

Activity-induced DPP-mediated VDF is uniquely observed in L-type (Ca_V_1.1-1.4) channels^13–17^, and contributes to various physiological processes^15,16,27,29,30,43,49,56^. However, the molecular origin and biophysical mechanisms underlying the VDF phenomenon remain elusive to date^13,14,17,32,57,58^. The results of this study provide the first evidence for the molecular endpoint of VDF by identifying a novel π-H and π-π quadrangle interaction, asymmetrically incorporated exclusively at the DI–DII PD interface of α_1_ subunits. Notably, replacing a single residue in Ca_V_2.1 channels with its Ca_V_1 counterpart converts the non-VDF-expressing Ca_V_2.1 channel into one that exhibits robust VDF. The reverse mutation in Ca_V_1.2 channels abolishes VDF, emphasizing the quadrangle’s critical role. Additionally, the complete inhibition of VDF by the common plastic leachable 2,4-DTBP, which disrupts π-H and π-π interactions at the DI–DII PD interfaces, underscores broader implications for voltage-gated ion channel pharmacology and pathophysiology.

Plastic leachables and microplastics are known to adversely affect Ca_V_ channels and underlying calcium signaling, contributing to various pathophysiologies, including endocrine dysfunction, cardiovascular complications, and neurodegenerative diseases such as autism and Alzheimer’s disease, among others^40,45,46,59,60^. However, the specific biophysical mechanisms targeted by these leachables remain poorly characterized. In this study, we identified complete inhibition of Ca_V_1.2 and Ca_V_1.3 L-type channel VDF by PP-stored solutions (***Fig. 1 and S3***) and the 2,4-DTBP leachable at nanomolar concentrations (***Fig. 4***)***, S7, S9, and S10***). Also, the ambiguity of L-type channel VDF inhibition or channels attaining a pre-facilitated state in PP and 2,4-DTBP solutions was ruled out by the lack of changes in Ca_V_1.2 V_a_ (***Fig. 2***)), current density (***Fig. 2 and S5***), and shift in open dwell time states (***Fig. 1 and 4***) in response to DPP. This finding is particularly striking because an excess concentration of 2,4-DTBP (18.3 ng/mL) has been observed in human urine samples^61^, and it has been regarded as an endocrine-disrupting chemical^62,63^. Given its lipophilicity, moderate molecular weight (∼206 Da), and low polar surface area, 2,4-DTBP is likely to favor passive diffusion across the blood-brain barrier and the placenta, thereby modulating L-type channel function in various tissue^62,64^. Indeed, exposure to 2,4-DTBP was found to increase mortality and induce anxiety-like behavior in zebrafish larvae by altering dopaminergic, serotonergic, and GABAergic signaling^65^; reduce pregnancy rates and impair kidney development in mice^62^; and impair osteogenic differentiation^46^ and adipogenesis^66^ in human mesenchymal stem cells, suggesting broad pathophysiological consequences.

Nonetheless, VDF inhibition by 2,4-DTBP enabled us to characterize the molecular endpoints and biophysical processes involved. A stable binding of 2,4-DTBP was observed at the DI–DII PD fenestration of α_1C_ and α_1D_ subunits by forming H-bonding with an asparagine residue in the DIIS6 helix (***Fig. 5***)***, S11, Video S1 and S2***). The binding pocket of 2,4-DTBP aligns with the recent discovery of a hydrophobic pocket that accommodates blocking lipids in this fenestration of Ca_V_1.2 and Ca_V_1.1 channel structures^5,67^. Importantly, N649 in rabbit Ca_V_1.1 (homologous to N741 in Ca_V_1.2 and N740 in Ca_V_1.3) was documented to exclusively engage in H-bond formation with verapamil’s tertiary amine group, a high-affinity L-type channel antagonist^6^. Modification of the tertiary amine was also shown to decrease its potency of L-type channel inhibition in smooth muscle cells^68^. Similarly, N741 in Ca_V_1.2 was found to engage in H-bonding with the anti-rheumatic drug tetrandrine^4^. Thus, 2,4-DTBP’s H-bonding with the conserved asparagine in the DIIS6 helix (***Fig. 5 and S11***) may broadly disrupt L-type channel pharmacology, including antihypertensive, anti-rheumatic, and angina pectoris therapies^2,6^. Importantly, the DIIS6 asparagine is conserved across all Ca_V_ and voltage-gated sodium (Na_V_) channels (***Fig. 7B and S14A*)**, suggesting an essential biophysical role. Additionally, the DI–DII PD interface

exhibits high sequence conservation, comprising hydrophobic residues across all Ca_V_ and Na_V_ channels that may support 2,4-DTBP binding^2,3,7,12^. Thus, this study highlights not only L-type channel VDF inhibition but also the broad-ranging implications of 2,4-DTBP on the physiological and pharmacological properties of Ca_V_ and Na_V_ channels.

This study importantly revealed an asymmetric π-H and π-π quadrangle interaction at the DI–DII PD interface that supports the DPP-induced VDF (***Fig. 5, 6, S11***, and ***S13***). In analogy, recent studies have also documented an asymmetric origin underlying the inactivation of Ca_V_ channels^39,69^. The reorientation of the amide group of the asparagine in DIIS6 by 2,4-DTBP indicates the need for stereospecificity in the quadrangle, which is supported by VDF inhibition in alanine or glutamine-substituted channels (***Fig. 5 and 6***). However, further structural studies will be required to confirm the reorientation of the π-H bond in the presence of 2,4-DTBP. Nonetheless, a complete rescue of VDF in the Ca_V_1.2-F737Y restoring the benzene ring geometry required for the stabilization of π-H and π-π quadrangle, but not in the Ca_V_1.2-F737A, argues well for stereospecificity (***Fig. 6***)). The stabilization of π-H and π-π quadrangle geometry facilitated the channel’s long open state, as evidenced by increased VDF and open probability/long-open dwell time of Ca_V_1.2 channels in response to DPP, but not in 2,4-DTBP-or PP-incubated solutions (***Fig. 1 and 4***). Supporting this, Ca_V_1.2-F737Y channels not only rescued VDF but exhibited a threefold increase in dwell time compared with Ca_V_1.2-WT, indicating a greater stabilization of the open state (***Fig. 4***)). This corroborates the proposed role of the DIIS6 phenylalanine in restricting inactivation of Ca_V_1.2 channels, in which small side-chain amino acids, such as alanine or glycine, displayed greater channel inactivation than bulkier substituents, such as phenylalanine^53^. Altogether, this suggests a crucial role of the π-H and π-π quadrangle geometry required for generating the DPP-mediated VDF in L-type channels (diagrammatically represented in ***Fig. 7F***)). However, conformational orientation of this π-H and π-π quadrangle in the resting and activated states requires further investigation, since all currently solved L-type channel structures represent the inactivated state^5,6,50–52^. Further structural/computational studies that reveal the resting, open, and intermediate transition states of L-type channels at varying depolarizing voltages will provide additional insight into the conformational rearrangements of the key π-H and π-π quadrangle interactions at the DI–DII PD interface.

Furthermore, Ca_V_2.1-T698F channels attaining L-type-like geometry for π-H and π-π quadrangle interactions exhibiting a substantial VDF in response to DPP (***Fig. 7C-7E***), highlight the central role of asymmetric π-H and π-π quadrangle interactions at the DI–DII PD interface in defining the molecular endpoint of VDF. This could be leveraged to dissect the physiological contributions of VDF to neuronal plasticity, given the widespread expression of L-type channels in the soma and postsynaptic structures^1,70,71^. Due to near-sequence conservation, the probable contribution of the DI–DII quadrangle to other forms of CDF and RGK-GTPase-mediated facilitation can also be hypothesized, which are observed in both Ca_V_1 and Ca_V_2 channels^18–24^.

In conclusion, these findings resolve a 30-year biophysical puzzle by revealing the π-H and π-π quadrangle interactions at the DI–DII PD interface as the molecular endpoint, along with their subtle conformational dynamics that drive the activity-dependent VDF of L-type channels with single-amino-acid precision.

## Methods

### Cell culture and transfection of Plasmid cDNAs

tsA-201 cells were routinely passaged and maintained in a humidified incubator with 5% CO_2_ and 95% air at 37°C in high-glucose containing Dulbecco’s Modified Eagle Medium (DMEM; 11965092, ThermoFisher) supplemented with 10% heat-inactivated Fetal Bovine Serum (FBS; A5256901, ThermoFisher) and 1% Penicillin-Streptomycin solution (ThermoFisher). Cells were passaged every 2 to 3 days after reaching ∼80-90% confluency. For electrophysiological recordings, cells were seeded onto 12-mm round coverslips placed in 35-mm dishes at 30-40% confluence. 20-24 hr after seeding, 2-3 μg of cDNAs of wild type (WD) or point mutated α□ subunit of L-type channel isoform Ca_V_1.2 (α_1C_, mouse, Addgene ID: 26572,^72^) or Ca_V_1.3 (α_1D_, rat, Addgene ID: 49332,^73^) in pcDNA6 backbone, was co-transfected with accessory subunits β_1b_ (rat) and α_2_δ_1_ (rat) in pcDNA3.1^53,74^ into tsA201 cells in varied combinations, specifically mentioned in the results section. The ratio of the plasmids used was α□, β_1b_, and α□δ□ at 1:0.75:0.5, respectively. In experiments involving P/Q-type currents, the cDNA of Ca_V_2.1 (α_1A_, human, Addgene ID: 140575^39^) was co-transfected with β_1b_ at a 1:0.75 ratio. 500 ng of eGFP in pcDNA3.1 was co-transfected with each sample to visualize transfected cells during whole-cell or single-channel recordings. In a subset of experiments, the Ca_V_1.2 cDNA was co-transfected with β_1b_, and C-terminal eGFP-tagged α□δ□ (α□δ□-eGFP) in ratios of 1:0.75:0.5 or 1:0.75:1. Unless otherwise mentioned, all experiments involving α□δ□ were performed using the untagged construct. All transfections in tsA-201 cells were performed using the established calcium-phosphate method^33^. 5-6 hours post-transfection, fresh DMEM supplemented with FBS was added, and the cells were maintained in a humidified incubator with 5% CO_2_ and 95% air at 30°C. All patch-clamp recordings were performed 36 to 72 hours post-transfection, and only eGFP-positive cells were selected for patching.

### Site-directed mutagenesis

Site-directed point mutations in the Ca_V_1.2 or Ca_V_2.1 cDNA were generated by long-range PCR using custom-designed forward and reverse primers that incorporated the mutated sequence. The forward and reverse primer sequences used are tabulated in **Table S2**. 2X Phanta Max Mastermix (P515, Vazyme, China) or 2X Fantom Master Mix (MM103, G2M, India) was used for PCR amplification of the desired mutants, following a 35-cycle reaction according to the manufacturer’s instructions. The amplified products were verified through agarose gel electrophoresis, followed by DpnI (R0176, NEB) restriction digestion and transformation into chemically competent TOP10 *E. coli* strains. Sanger’s cDNA sequencing was done to verify the mutations at the precise location of the plasmid.

### Hippocampal pyramidal neuron culture

Dissociated cultures of rat hippocampal pyramidal neurons were prepared from P0–P1 rat pups following all the institutional ethics protocols. After anesthetizing on ice, brains were surgically dissected, and both hippocampi were isolated in ice-cold Hibernate media, using a stereo zoom microscope (Magnus, India). Neurons were dissociated with 0.25% trypsin solution at 37°C for 10 minutes, followed by washing and mechanical trituration in Hibernate medium supplemented with 1% B-27, 2% Glutamax, and 1% Normocin (InvivoGen). The single-cell suspension was then centrifuged and resuspended in Neurobasal-A medium supplemented with 2% B-27, 2% Glutamax, and 1% Normocin. The neuronal suspension was plated on 0.1% poly-L-lysine-coated 12 mm round glass coverslips, and cells were grown on coverslips in neuronal growth medium (NGM) composed of Neurobasal-A, 2% Glutamax, 2% B-27 supplement, and 1% Normocin at 37°C in a 5% CO_2_ and 95% air humidified incubator^33,75^. Post-7-8 days in vitro (DIV) neuronal cultures were used for patch-clamp experiments. All media components were obtained from ThermoFisher, unless otherwise specified.

### Whole cell patch clamp electrophysiology

Whole-cell patch clamp recordings were performed on transfected tsA-201 cells or cultured hippocampal neurons maintained at room temperature (22-24), using a Multiclamp 700B amplifier run on Digidata 1440B and pClamp 11.3 software (Molecular Devices, USA). All whole-cell current signals were acquired using a 2 kHz analog Bessel filter and digitized at 20 kHz. After achieving whole-cell configuration, the whole-cell capacitance was neutralized, followed by 70% series resistance compensation and online leak subtraction using the Commander module of the Multiclamp 700B. For isolating Ca_V_1.2 currents, transfected tsA-201 cells were bathed in an external solution containing (in mM) 110 NaCl, 10 TEA-Cl, 1 MgCl_2_, 5 BaCl_2_, 10 HEPES, and 10 glucose, adjusted to pH 7.4 with NaOH. Ca_V_1.3 currents in transfected tsA-201 cells were recorded in bath solutions composed of (in mM): 110 NaCl, 10 CsCl, 5 BaCl_2_, 1 MgCl_2_, 10 mM HEPES, and 10 glucose, with the pH adjusted to 7.4 with NaOH.Voltage-gated calcium currents in the primary cultured hippocampal neurons were measured in external solutions containing (in mM) 90 CsCl, 40 TEA-Cl, 5 BaCl_2_, 1 MgCl_2_, 10 HEPES, 20 Glucose adjusted to pH 7.4 with CsOH. Thick-walled glass pipettes (with O.D., 1.5 mm and I.D., 0.86 mm) containing filament made of borosilicate (Sutter Instruments, USA) are used for patch clamping with an intracellular solution consisting of (in mM) 110 CsCl, 10 TAE-Cl, 10 HEPES, 1 MgCl_2_, and 10 EGTA, with the pH adjusted to 7.3 with CsOH, attaining a resistance of 4-6 MΩ.

The current-voltage (I-V) relationships were obtained by applying 500-msec voltage steps from −80 mV to +60 mV in 10-mV increments, and peak inward currents were measured at each voltage. I–V curves were fitted using the built-in Boltzmann I–V function in the Origin software.

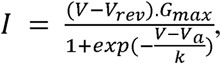

Where I is the peak current, at voltage V, V_rev_ is the reversal potential, G_max_ is the maximum conductance, V_a_ is the half-activation voltage, and k is the slope factor. Similarly, the conductance of Ca_V_1.2 and Ca_V_1.3 channels at specific voltages was determined using

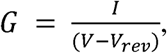

In which the V_rev_ was obtained from the fitted I-V plot of the same channel under investigation. The conductance-voltage (G-V) relationship was calculated by fitting the G/G_max_ values with the equation:

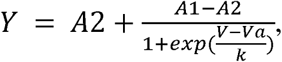

Where Y = G/G_max_, A1 and A2 are the maximum and minimum values of the G/G_max,_ respectively. The residual current fraction after 400 ms, R_400_, was calculated as the ratio of the current amplitude measured 400 ms after the onset of the voltage step to the corresponding peak current amplitude.

Voltage-dependent facilitation (VDF) was assessed using a paired-pulse protocol, in which cells were held at -80 mV, and a test pulse (referred to as P1) of 0 mV for Ca_V_1.2 or -10 mV for Ca_V_1.3 lasting 20 msec was applied to elicit the baseline current level at the start. Following a 25-second recovery, a depolarized prepulse (DPP) ranging from 0 mV to 180 mV in increments of 20 mV was stimulated for 25 msec. After 5 msec of DPP, a second test pulse (referred to as P2) of either 0 mV for Ca_V_1.2 or -10 mV for Ca_V_1.3 for 20 msec was applied to determine the magnitude of facilitation. Peak current amplitudes obtained with or without DPPs were used to calculate the % of VDF using the following formula:

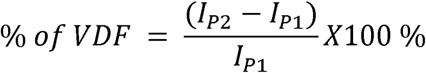

Where I_P2_ refers to the peak current obtained by the test pulse applied after the DPP, while I_P1_ refers to the basal peak current recorded prior to the DPP stimulus.

To examine the DPP-induced VDF and channel activation properties, a 25 ms DPP of +140 mV was applied before a 500 ms test pulse ranging from −80 to +60 mV in 10 mV increments, with a 10 ms interval between the DPP and the test pulse. In case of paired recordings, the VDF of Ca_V_1.2 channels was assessed with and without a +140 mV DPP, following a 0 mV test pulse, by exchanging the bath solutions from glass to PP-incubated solutions or to solutions containing 500 nM 2,4-DTBP.

To determine the effect of specific plastic leachables on the L-type channel VDF, cells were incubated with the respective leachable at varying concentrations (as mentioned in the results section) for at least 7-10 minutes prior to initiation of VDF measurements. The maximum VDF in response to a series of DPP voltages ranging from 0-180 mV was determined by fitting the percentage of facilitation plots with the Richards growth equation in Origin:

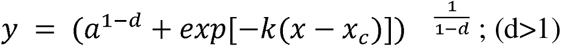

Where “a” represents the asymptotic maximum facilitation, and k is the growth rate constant.

All electrophysiological data analysis was performed using ClampFit (Molecular Devices, USA) in pClamp 11.3, unless otherwise specified. For data representation, capacitance transients were hidden from all the current recording traces. All patch-clamp chemicals were obtained from Sigma-Aldrich, unless otherwise specified.

### Single-channel electrophysiology

Cell-attached single channel recordings were performed on tsA-201 cells coexpressing Ca_V_1.2 (mouse) WT or point-mutated channels along with the β_1b_ (rat), at room temperature (22-24). The single-channel currents were acquired using the Multiclamp 700B amplifier with a 5 GΩ feedback resistance. All single-channel signals were acquired using a 1.2 kHz basal filter and digitized at 7.14 kHz with the Digidata 1440B and pClamp 11.3 software. Cells were bathed in an external solution containing (in mM) 140 K Aspartate, 5 EGTA, 10 HEPES, 1 MgCl_2_, and 10 Glucose at pH 7.4 adjusted with KOH, to attain a membrane potential of 0 mV. Thick-walled glass pipettes (as mentioned before) having resistances of 6-8 MΩ, filled with pipette solution consisting of (in mM) 110 BaCl_2_, 10 HEPES, and 5 TEA-Cl, pH adjusted to 7.4 with Ba(OH)_2_. The pipette solution contained 1 μM of (S)-(−)-BAY-K-8644 (19988, Cayman Chemical) to selectively increase the L-type channel activity. The DPP-induced gating transition of the Ca_V_1.2 channel was reported to be distinct from the gating event of the Bay-K-8644 allosteric agonist^32^, which also shifts the channel to a higher open probability state, but via a different mechanism^6^. This facilitates the examination of L-type channel VDF in the presence of Bay-K-8644. After formation of the gigaseal, cells were continuously held at -80 mV, and the respective channel activity was recorded by stepping to 0 mV for 2 sec. For eliciting VDF, a 100 mV DPP of 100 msec was applied, followed by a second 0 mV step of 2sec. To study the effect of PP-stored solutions or 2,4-DTBP on L-type channel VDF, cells were exposed to a bath solution containing an equivalent volume of PP-tube stored DD water, matching the conditions used in whole-cell experiments. Cells were left in the solution for at least 5 minutes before data acquisition. For experiments involving 2,4-DTBP, a concentration of 500 nM was used in the bath solutions for at least 5 min prior to cell-attached patch-clamp recordings. The open probability of single-channel traces was calculated using Clampfit software in pClamp11. Single-channel current amplitude and open dwell-time events were extracted using the pyabf python package. Subsequently, the single-channel amplitude values were fitted with a two-peak Gaussian histogram function in Origin-8 to calculate the mean single-channel amplitudes. For open dwell time analysis, opening events resulting from a single channel were considered, while multiple-opening events were excluded. A logarithmic histogram method was used to plot the open dwell time distributions^76^, with a constant bin width of 0.2 ln units after transforming dwell times to ln(t). Open-time distributions were fitted using a non-linear least-squares algorithm following previously reported methodology^77–79^. The following equation was used to fit the logarithmic open-dwell time distributions with a sum of exponential components,

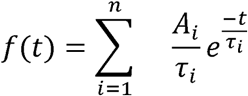

where A_i_ and τ_i_ are the fractional amplitudes and time constants of the components, respectively.

### Extraction of plastic leachables

To evaluate leaching behaviour, aqueous solutions of 1M barium chloride (BaCl) (B0750, Sigma Aldrich) or double-distilled (DD) water were stored in PP tubes for 7-14 days in 50 mL Falcon Polypropylene (PP) Centrifuge Tubes (352070, Corning Life Sciences; 339652, ThermoScientific; 546041, Tarsons) that are commonly used in laboratories. As a control, identical 1 M BaCl or DD water samples were incubated in borosilicate laboratory glass bottles (B1501021, Borosil). For characterization of leachables, the aqueous BaCl or DD water was isolated by liquid-liquid extraction. Briefly, 50 mL of the PP-or glass-bottle-incubated solution was transferred to a separating funnel, and HPLC-grade ethyl acetate (Catalog no. E/0906/17, Fisher Chemical) was added. The extraction was repeated three times, and the combined organic fractions were collected and concentrated to 1 mL using a rotary evaporator (Buchi, IKA). Afterward, the samples were further examined to detect specific leachables.

### Thin Layer Chromatography (TLC)

Concentrated extracts of 1M BaCl stored in PP tubes for 30-40 days were run on TLC plates (Merck Millipore) using a mobile phase of ethyl acetate: hexane (1:9, v/v). After development, TLC plates were examined under ultraviolet light at 254 nm and 365 nm, and spots corresponding to leachables were documented. Preparative TLC was used to collect fractions of interest, in which concentrated extracts were applied in multiple layers to large sized TLC plates, with intermittent drying. Plates were developed in the same mobile phase and examined under UV light. Bands corresponding to leachables were marked, scraped, and extracted in Ethyl acetate. The samples were sonicated, then vortexed and centrifuged at 10,000 rpm for 10 minutes. The supernatant was further concentrated using a rotary evaporator. The resulting extracts were evaluated using Gas Chromatography-Mass Spectrometry (GC-MS) or Liquid Chromatography-Mass Spectrometry (LC-MS) to characterize leachables.

### GC-MS and LC-MS

Concentrated samples of liquid-liquid extracted 1M BaCl in ethylacetate were further diluted and analyzed by GC-MS using the Agilent 7000D Triple Quadrupole GC/MS system. The spectra were compared with those of extracts prepared from control samples of identical 1 M BaCl stored in glass bottles for similar time durations. Similarly, an Agilent 1260 Infinity LC system, coupled with a Quadrupole Time-of-Flight (QTOF) mass spectrometer, was used to analyze TLC extracts in chloroform by LC-MS. The distinct peaks observed in GC-MS or LC-MS sample extracts, compared to control extracts, were referenced in the National Institute of Standards and Technology (NIST) mass spectral database for tentative compound identification. The extracted ion chromatogram (EIC) of tentative compounds was further compared with the EIC patterns of commercial standards to characterize and validate them by performing similar GC-MS and LC-MS analyses in ethyl acetate and methanol, respectively.

### Docking and Molecular Dynamics (MD) Simulations

A blind docking was employed to identify the potential binding site of the 2,4-DTBP (PubChem) within the L-type calcium channel Ca_V_1.2 (PDB:8WE6)^50^ and Ca_V_1.3 (PDB:7UHG)^51^ protein complex using AutoDock4.2. The Ca_V_1.2 structural complex contained human α_1C_, β_3_, and α_2_δ_1_ subunits. Similarly, the Ca_V_1.3 structural complex contained human α_1D_, β_3_, and α_2_δ_1_ subunits. From the thousand generated poses, the best binding configuration of 2,4-DTBP with its respective channels was selected based on the lowest binding energy and the maximum number of possible interactions. To reduce computational costs and improve performance, and given the nature of the binding site, the voltage-sensing domains (VSDs) were truncated, retaining only the pore domains of Ca_V_1.2 and Ca_V_1.3 for all-atom molecular dynamics (MD) simulations in both ligand-bound and ligand-free states. All simulation trials were extended to 500 ns to ensure adequate sampling of protein-ligand interactions and to evaluate the stability of the bound ligand.

The selected protein and ligand complex of 2,4-DTBP with the α_1C_ subunit of Ca_V_1.2 or the α_1D_ subunit of Ca_V_1.3, obtained from molecular docking, was embedded inside the lipidic membrane environment consisting of 1-Palmitoyl-2-oleoyl-sn-glycero-3-phosphocholine (POPC) using CharmGui membrane builder web server^80^. The entire system was then solvated with transferable intermolecular potential 3-point (TIP3) water molecules, followed by charge neutralization in the presence of 150 mM NaCl and 20 mM BaCl_2_. The CHARMM36m force field was used to run the MD simulations. To ensure independent sampling and assess the reproducibility of the results, multiple simulation replicas were generated with different random seeds, providing unique initial velocities and distinct starting trajectories for each simulation.

The systems were energy-minimized using the steepest descent function in GROMACS 2020.4 to remove steric clashes and relax the initial configuration. Afterward, positional restraints were applied to the protein backbone, side chains, and lipid head groups. Long-range electrostatics were treated using the Particle Mesh Ewald (PME) algorithm, while the LINCS algorithm was used to constrain the covalent bond involving H atoms. A multistep equilibration process was employed to gradually relax the systems while preserving the protein-2,4-DTBP complex and transmembrane integrity. An initial equilibration was performed in two successive steps, each lasting 1.25 ns with a 1-femtosecond (fs) integration time step. Positional restraints were applied to the protein backbone, side chains, and lipid headgroups in these steps, and the temperature was held at 303.15K using the Berendsen Thermostat. Subsequent equilibration steps were performed using semi-isotropic pressure coupling with Berendsen Barostat. From the fourth equilibration step onwards, the positional and dihedral restraints were progressively weakened, and the integration time step was 2 fs. The final production runs were 500 ns for Ca_V_1.2 and Ca_V_1.3, with all restraints removed. The trajectory coordinates, velocities, and forces were saved every 100 ps for the post-production structural and dynamic analysis. The Python package MDAnalysis and PyMOL were used to analyse the saved trajectories. The centroid-to-centroid distance between nearby residues was calculated using MDAnalysis. The RMSD of the protein backbone was determined using trajectories in which the disordered extracellular regions of the PD were excluded from the analysis. Hydrogen bond occupancy between N741/N740 and 2,4-DTBP was analyzed using VMD. Only polar atoms (N, O, S, and F) were considered as potential hydrogen-bond donors and acceptors, with a donor-acceptor distance cutoff of 3.5 Å and an angle cutoff of 40°. A Principal Component Analysis (PCA) was used to capture the motion of the F737 and N741 sidechains in Ca_V_1.2, as well as sidechains of F736 and N740 in Ca_V_1.3, by selecting non-backbone atoms using MDAnalysis. The simulation trajectories were structurally aligned to a common reference frame to remove the overall rotation and translation. To capture the local conformational changes, PCA was performed on the complete set of side-chain atomic coordinates rather than on a single distance-based parameter. PCA was performed on the atomic covariance matrix, and the trajectories were projected onto the first two principal components (PCs, PC1 and PC2), which captured the largest variance in side-chain motion. To identify the discrete orientational states of the side chains, the PCA-derived PC1 and PC2 values were clustered using the scikit-learn Python package’s k-means algorithm. Based on the separation and variance of the data, the conformational ensembles were clustered into distinct groups, representing different side-chain orientations. PyMOL and VMD were used to visualize trajectories and to prepare graphical illustrations for the figure.

### Homology Modeling

To visualize the structural effects of the mutations within the quadrangle region, homology-based models of the channels were generated using the SWISS-MODEL server. The experimentally determined cryo-EM structure of Ca_V_1.2 (PDB ID: 8WE6) was used as the template for model construction. The generated structures were subsequently used for comparative structural analysis and their potential impact on quadrangle geometry.

### Multiple Sequence Alignment

Protein sequences analyzed in this study were retrieved from the UniProt database using their corresponding accession numbers, which are presented in Table S3. Representative voltage-gated calcium channel α_1_ subunit sequences from different species [h(human), m(mouse), r(rat), rb(rabbit)] were selected to enable comparative sequence analysis. Multiple sequence alignment (MSA) was performed using the Clustal Omega web server with default parameters to identify conserved and divergent regions across the selected calcium channel isoforms. The resulting alignments were used for subsequent comparative analyses.

### Statistical analysis

All quantitative data are presented as mean ± SEM, unless otherwise stated. An unpaired Student’s t-test statistical analysis was performed to compare the difference between two sets of independent data. One-way ANOVA was used to compare multiple groups with WT or control, unless otherwise explicitly mentioned. An unpaired Student’s t-test was used to assess the DPP-induced change in half maximal channel activation, V_a_. To compare DPP-induced VDF across a range of depolarizing voltages (0–180 mV), a two-way ANOVA was used. In the case of paired solution exchange experiments, a paired non-parametric Student’s t-test was used to determine significance between data sets. Single-channel parameters, including current amplitude and open probability, were analyzed using a non-parametric paired Student’s t-test, unless otherwise mentioned. One-way ANOVA with pairwise comparisons was used to assess the significance of differences among the measured open-dwell-time states. All results were considered statistically significant at P < 0.05 or less.

## Supporting information

Cav1.2 with 2,4-DTBP

Cav1.3 with 2,4-DTBP

## Data Availability

All raw datasets generated and custom-written Python codes used for analysis are available in the Zenodo data repository with DOI: **10.5281/zenodo.22869244**. The source data is provided in Excel sheets along with the manuscript.

## Acknowledgements

The authors acknowledge the Beagle MBU computational cluster and SERC data center, located at the Indian Institute of Science (IISc), Bangalore, for providing kind access. This work is funded by the Indian Council of Medical Research (ICMR) (grand IDs: IIRP-2023-0253, FIW-2024-01-0000000061, and IIRPSG-2025-01-01202) (G.S.), Department of Biotechnology (grant ID: BT/PR47597/BMS/85/46/2024) (G.S.), startup research support, IISc Bangalore (G.S.), student fellowship, IISc (A.H.), and funding support from the Ministry of Education (MoE), Govt. of India. G.W.Z. is supported by a discovery grant from NSERC.

## Author Contributions

A.H. generated point mutagenesis cDNA clones and performed all the electrophysiological, docking, and MD simulation experiments. V.T.K. and V.G.E. conducted the leachable characterization studies. A.H., V.T.K., V.G.E., A.K. and G.S. analyzed the results. A.H., V.T.K., and G.S. wrote the first draft of the manuscript, followed by input and suggestions from all the authors. G.S. and A.K. supervised the leachable identification studies. G.W.Z., A.K., and G.S. provided the necessary reagents and supplies, as well as valuable insights. G.S. conceptualized and supervised the overall study.

## Competing Interests

The authors declare no competing interests

## Supplementary Figure legends

**Figure S1:**
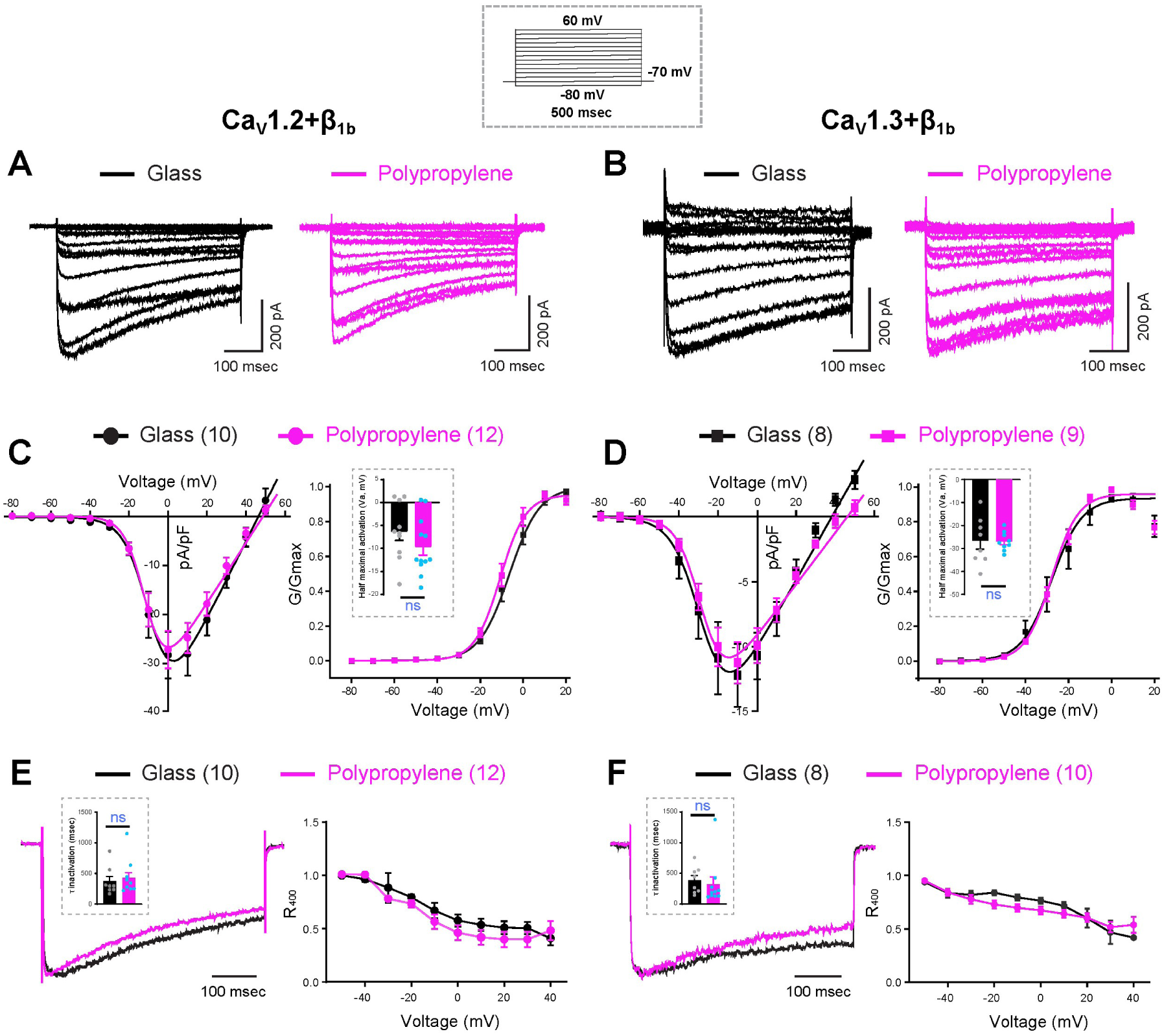
Buffer solutions stored in PP tubes do not alter activation and inactivation properties of Ca_V_1.2 and Ca_V_1.3 channels coexpressed with β_1b_ subunit. **(A and B)** Representative whole-cell current traces of Ca_V_1.2 **(A)** and Ca_V_1.3 **(B)** channels coexpressed with β_1b_ subunit in tsA-201 cells recorded in external solutions stored in glass bottles (black) or PP tubes (magenta). The voltage protocol is shown in the dotted box above. **(C and D)** Current density (pA/pF) and normalized conductance (G/G_max_) versus voltage plots of Ca_V_1.2 **(C)** and Ca_V_1.3 **(D)** show no significant differences when recorded in solutions stored in glass bottles (black) and PP tubes (magenta). Insets show half-maximal activation voltages. **(E and F)** Normalized current traces of Ca_V_1.2 **(E)** and Ca_V_1.3 **(F)** at 0 mV show channel inactivation in glass (black) or PP tubes (magenta). Insets display time constants of inactivation (τ) from exponential fits of 0 mV traces. R_400_ plots represent the ratio of residual current at 400 msec to the peak current amplitude across voltage steps ranging from -50 to +40 mV. All plots represent mean ± SEM. Experimental replicate cell numbers are indicated in brackets. The unpaired Student’s t-test is used for statistical significance. ns, non-significant.

**Figure S2:**
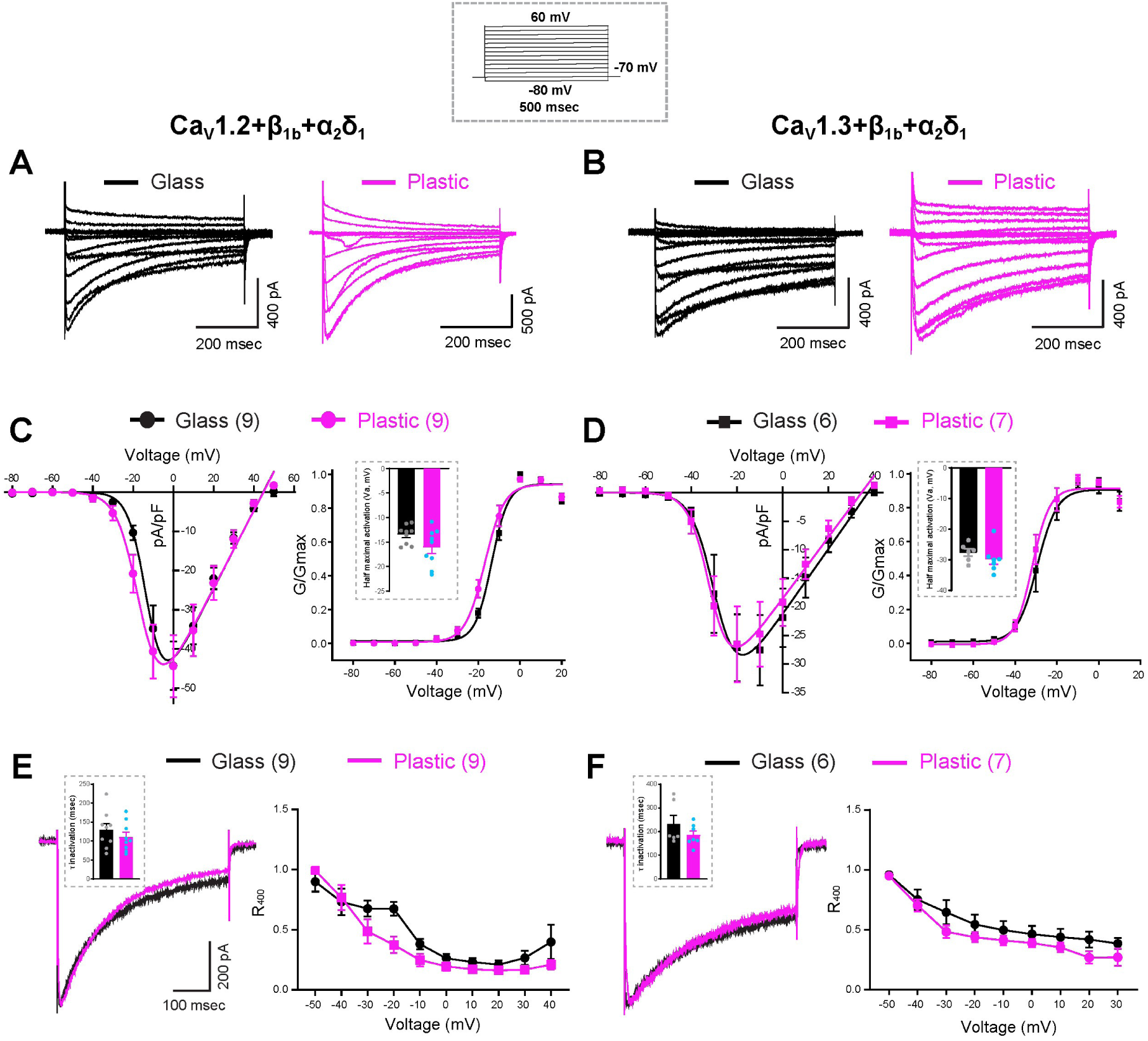
Buffer solutions stored in PP tubes do not alter activation and inactivation properties of Ca_V_1.2 and Ca_V_1.3 channels coexpressed with β_1b_ and α_2_δ_1_subunits. **(A and B)** Representative whole-cell current traces of Ca_V_1.2 **(A)** and Ca_V_1.3 **(B)** channels coexpressed with β_1b_ and α_2_δ_1_subunit in tsA-201 cells recorded in external solutions stored in glass bottles (black) or PP tubes (magenta). The voltage protocol is shown in the dotted box above. **(C and D)** Current density (pA/pF) and normalized conductance (G/G_max_) versus voltage plots of Ca_V_1.2 **(C)** and Ca_V_1.3 **(D)** show no significant differences when recorded in solutions stored in glass bottles (black) and PP tubes (magenta). Insets show half-maximal activation voltages. **(E and F)** Normalized current traces of Ca_V_1.2 **(E)** and Ca_V_1.3 **(F)** at 0 mV show channel inactivation in glass (black) or PP tubes (magenta). Insets display time constant of inactivation (τ) from exponential fits of 0 mV traces. R_400_ plots represent the ratio of residual current at 400 msec to the peak current amplitude across voltage steps ranging from -50 to +40 mV. All plots represent mean ± SEM. Experimental replicate cell numbers are indicated in brackets. Unpaired Student’s t-test used for statistical significance. ns, non-significant.

**Figure S3:**
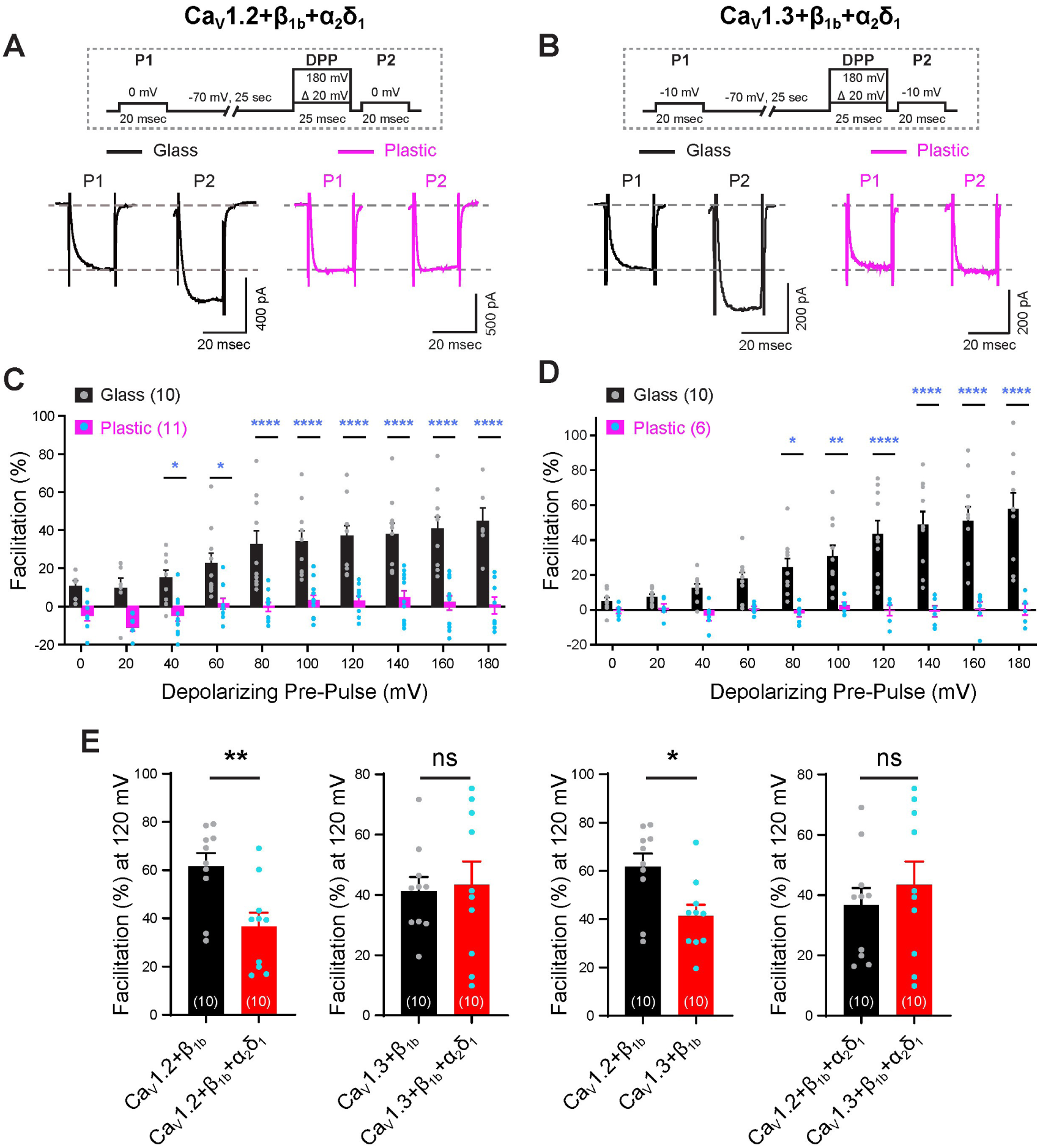
Buffer solutions stored in PP tubes potently inhibit the VDF of Ca_V_1.2 and Ca_V_1.3 L-type channels coexpressed with β_1b_and α_2_δ_1_ subunits: **(A and B)** Representative whole-cell current traces of L-type Ca_V_1.2 **(A)** and Ca_V_1.3 **(B)** channels coexpressed with β_1b_ and α_2_δ_1_subunits in tsA-201 cells recorded in external solutions stored in glass bottles (black) or PP tubes (magenta). The voltage protocol is shown in the dotted box above. **(C and D)** Bar plot quantifications depict percentage facilitation of Ca_V_1.2 **(C)** and Ca_V_1.3 **(D)** whole-cell currents, measured as peak current difference between P2 and P1 voltage steps (P2-P1), in solutions stored in glass bottles (black) or PP tubes (magenta). Cell numbers represented in brackets. Two-way ANOVA is used for statistical significance. **(E)** Bar plots depict percentage VDF (P2 over P1 at 0 mV after 120 mV DPP) for indicated L-type channel combinations. Unpaired Student’s t-test used for statistical significance. All plots represent mean ± SEM. Experimental replicate cell numbers are indicated in brackets. *p ≤ 0.05, **p ≤ 0.01, ***p ≤ 0.001, ****p ≤ 0.0001; ns, non-significant.

**Figure S4.**
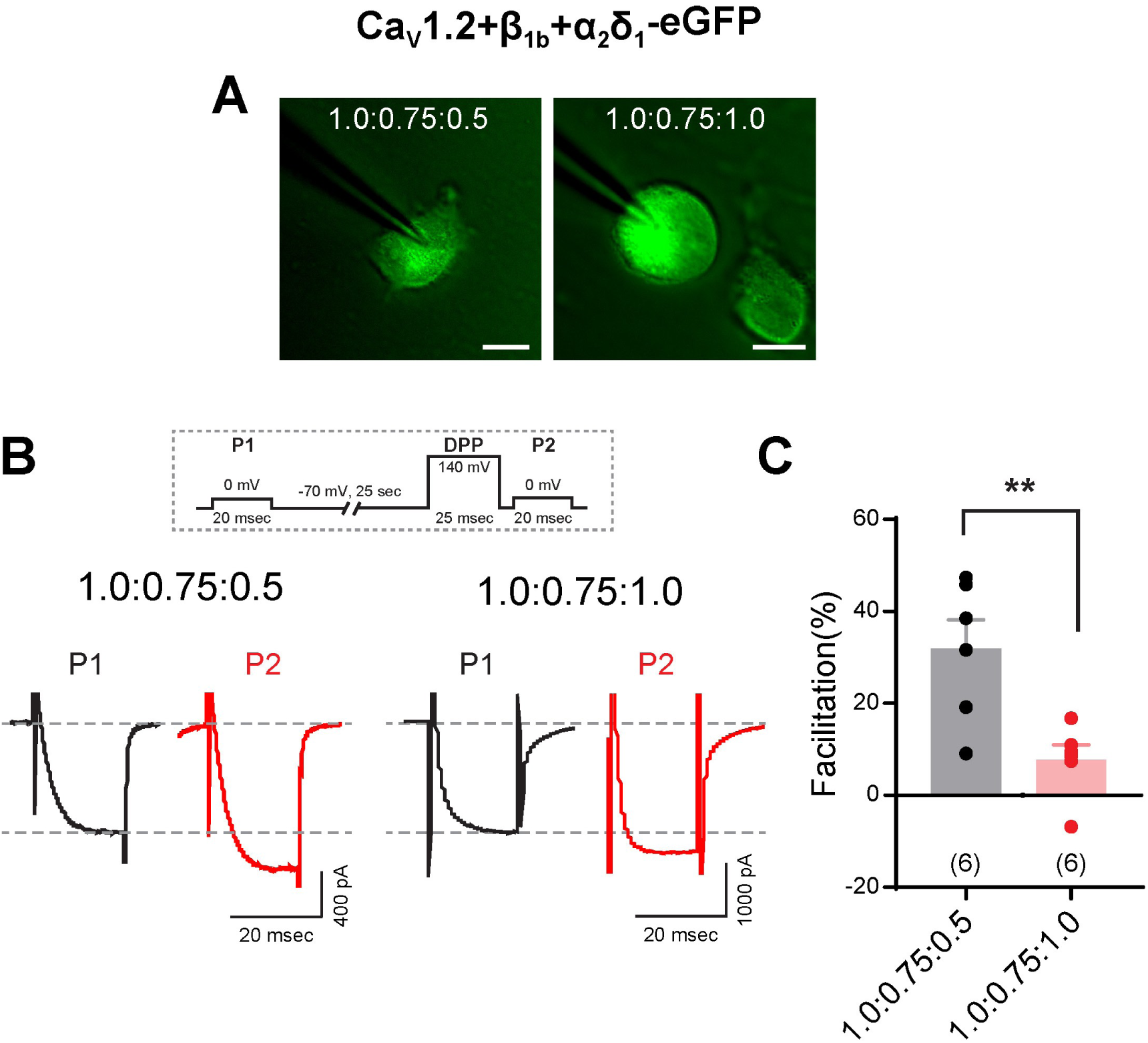
α_2_δ_1_ expression interferes with DPP-induced VDF of Ca_V_1.2 channels. **(A)** Representative fluorescence images of tsA-201 cells coexpressing Ca_V_1.2, β_1b_, and α_2_δ_1_-eGFP at transfection ratios of 1:0.75:0.5 and 1:0.75:1.0, respectively. Scale bars, 10 μm. **(B)** Representative whole-cell Ba² current traces of Ca_V_1.2 coexpressed with β_1b_ and α_2_δ_1_in specified ratios at 0 mV, in external solutions stored in glass bottles before (black) and after (red) a DPP of +140 mV. The voltage protocol used is shown in the dotted box above. **(C)** Bar plot quantification depicts the percentage of Ca_V_1.2 VDF coexpressed with β_1b_ and α_2_δ_1_ in specified ratios, following a DPP to +140 mV. Bar plots represent mean ± SEM. Experimental repeat cell numbers are indicated in brackets. An unpaired Student’s t-test is used for determining statistical significance. **p ≤ 0.01.

**Figure S5.**
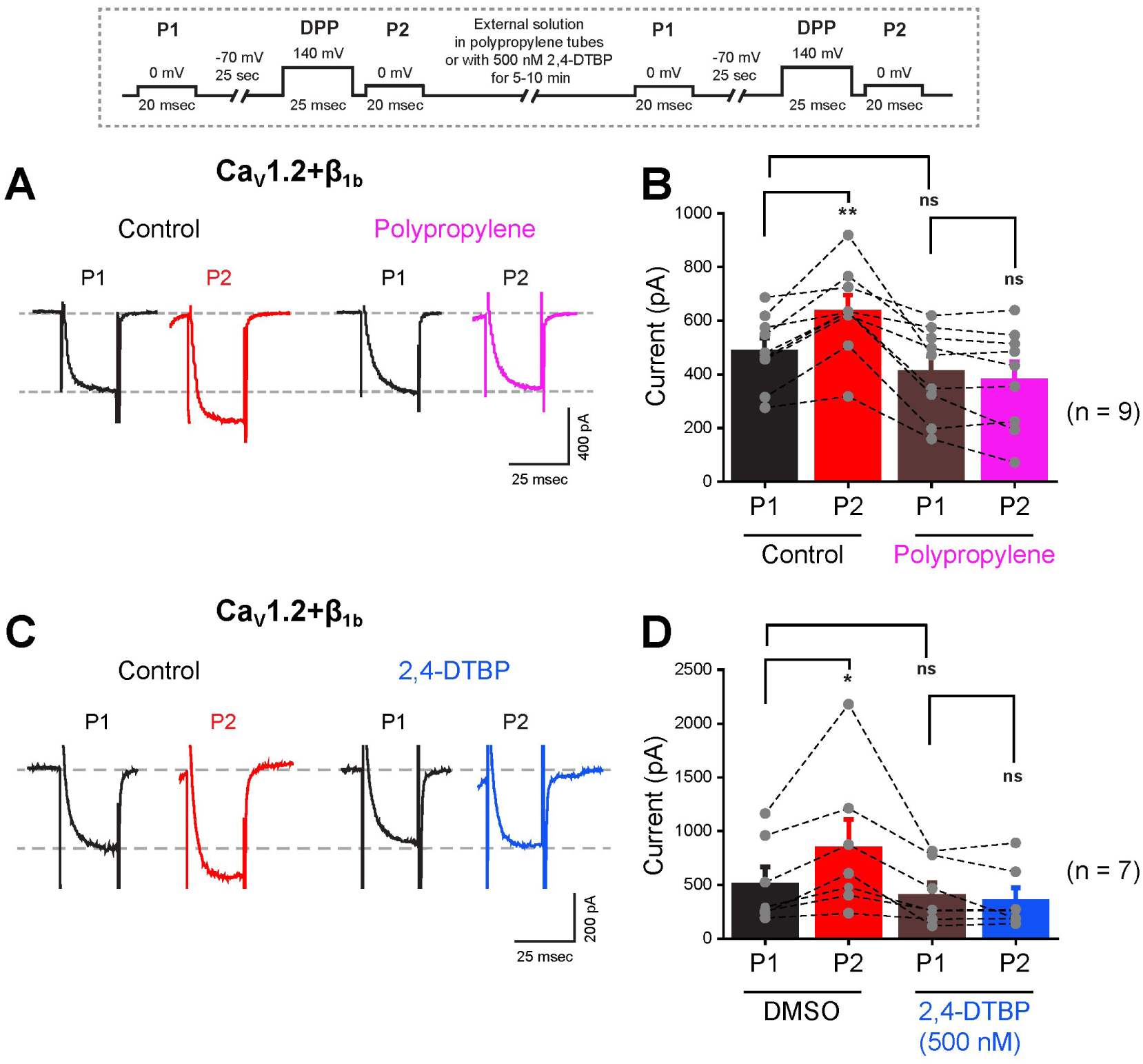
PP-stored solution and 2,4-DTBP abolish DPP-induced VDF of Ca_V_1.2. **(A)** Representative whole-cell Ba² current traces of Ca_V_1.2 coexpressed with β_1b_ at 0 mV before and after a DPP of +140 mV in external solutions stored in glass bottles (control), followed by 5-10 incubation in PP-stored solutions. **(B)** Bar plot quantification of Ca_V_1.2 VDF in glass bottle (control) and PP incubated solutions. **(C)** Representative whole-cell Ba² current traces of Ca_V_1.2 coexpressed with β_1b_ at 0 mV before and after a DPP of +140 mV in solutions containing DMSO, followed by 5-10 min incubation in 500 nM 2,4-DTBP. **(D)** Bar plot quantification of Ca_V_1.2 VDF in DMSO (control) and 500 nM-2,4-DTBP containing solutions. The voltage protocol is shown in the dotted box above. Bar plots represent mean ± SEM. Experimental repeat cell numbers are indicated in brackets. One-way ANOVA is used for determining statistical significance. *p ≤ 0.05, **p ≤ 0.01, ***p ≤ 0.001, ****p ≤ 0.0001; ns, non-significant.

**Figure S6:**
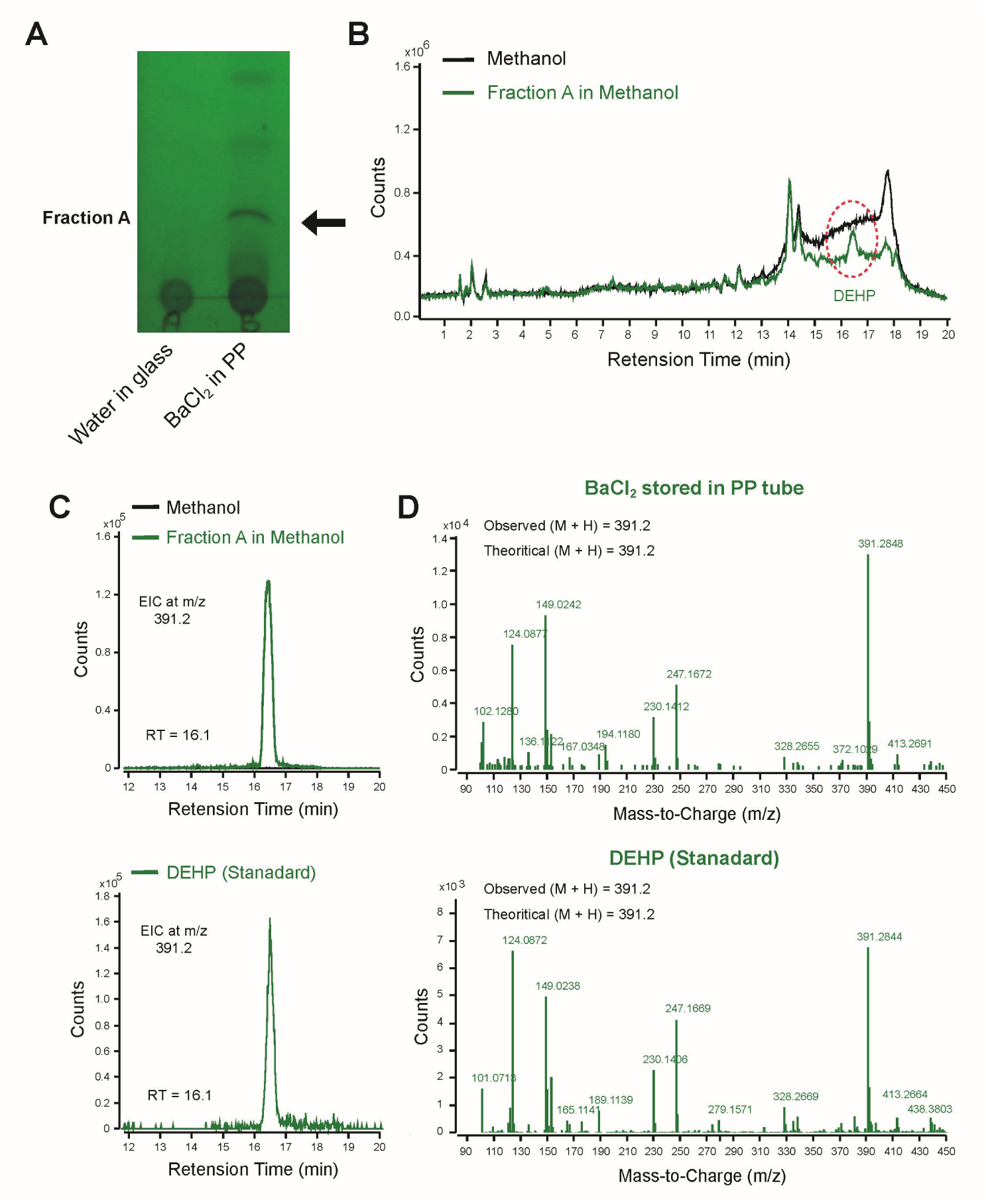
LC-MS analysis confirms DEHP leaching from aqueous BaCl_2_ stored in PP tubes: **(A)** TLC of extracts from 1 M aqueous BaCl_2_ stored in PP tubes vs. water in glass bottles. Distinctive fraction-A band marked with an arrow. Ethyl acetate and hexane in 1:9 ratio as the mobile phase. **(B)** LC-MS comparative chromatogram of fraction-A (methanol extract) vs methanol solvent control. The distinctive fraction-A peak is circled in the chromatogram. **(C)** Extracted ion chromatogram (EIC) at 391.2845 m/z for fraction-A and DEHP commercial standard displaying a retention time of 16.148 min and 16.154 min, respectively. **(D)** Mass spectrum at 391.2845 m/z for fraction-A and DEHP standard, showing theoretical and observed molecular masses.

**Figure S7:**
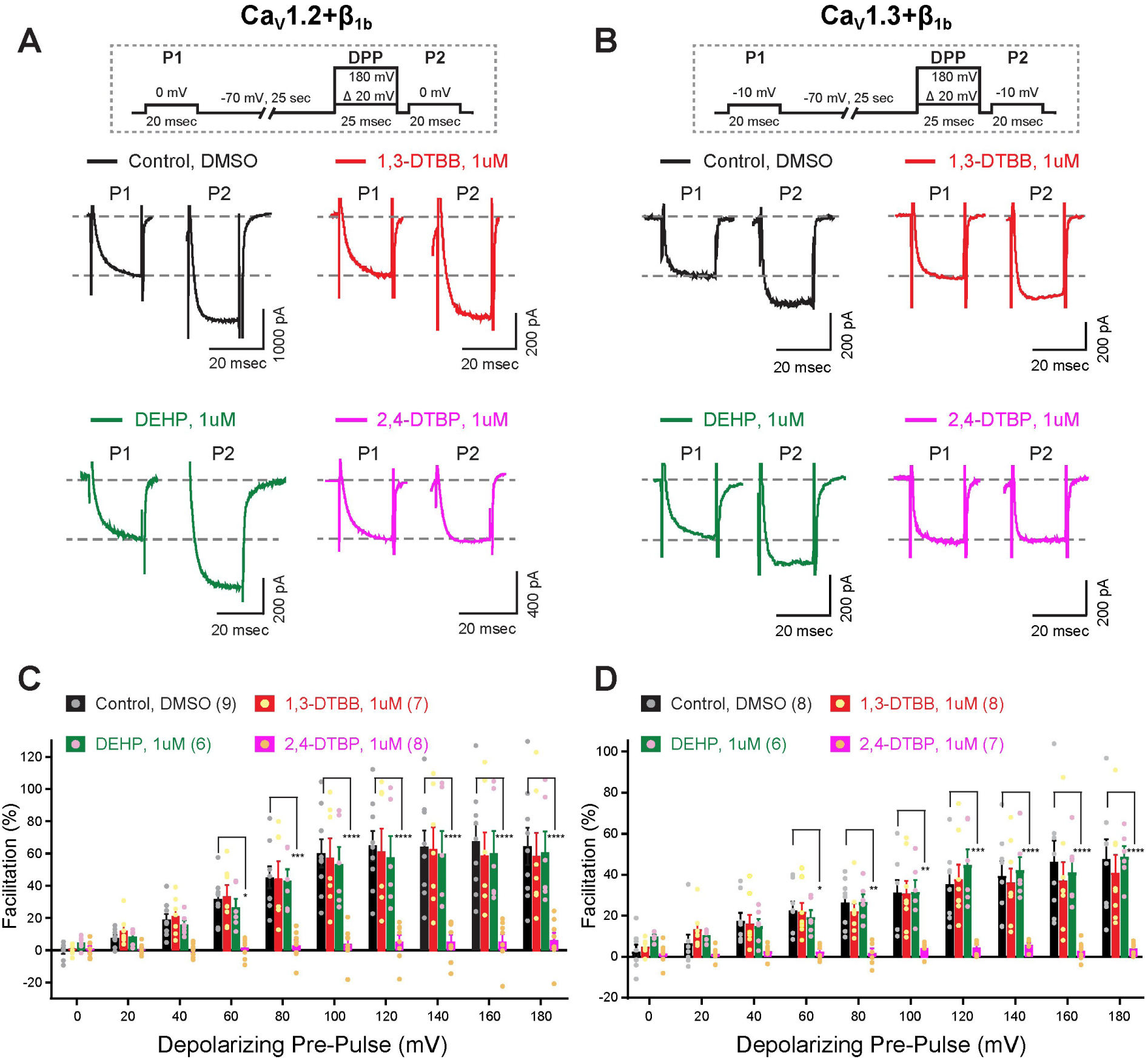
2,4-DTBP selectively inhibits Ca_V_1.2 and Ca_V_1.3 channel VDF: **(A and B)** Representative whole-cell current traces of L-type Ca_V_1.2 **(A)** and Ca_V_1.3 **(B)** channels coexpressed with β_1b_ subunit in tsA-201 cells recorded in external solutions containing 1 μM 1,3-DTBB (red), DEHP (green), or 2,4-DTBP (magenta), or DMSO (black). The voltage protocol is shown in the dotted box above. **(C and D)** Bar plot quantifications depict VDF percentage of Ca_V_1.2 **(C)** and Ca_V_1.3 **(D)** whole-cell currents, measured by the difference between peak current amplitudes of P2 and P1 (P2-P1) voltage steps, in external solutions containing 1,3-DTBB (red), DEHP (green), or 2,4-DTBP (magenta), or DMSO (black). Two-way ANOVA is used for statistical significance. All plots represent mean ± SEM. Experimental replicate cell numbers are indicated in brackets. *p ≤ 0.05, **p ≤ 0.01, ***p ≤ 0.001, ****p ≤ 0.0001.

**Figure S8:**
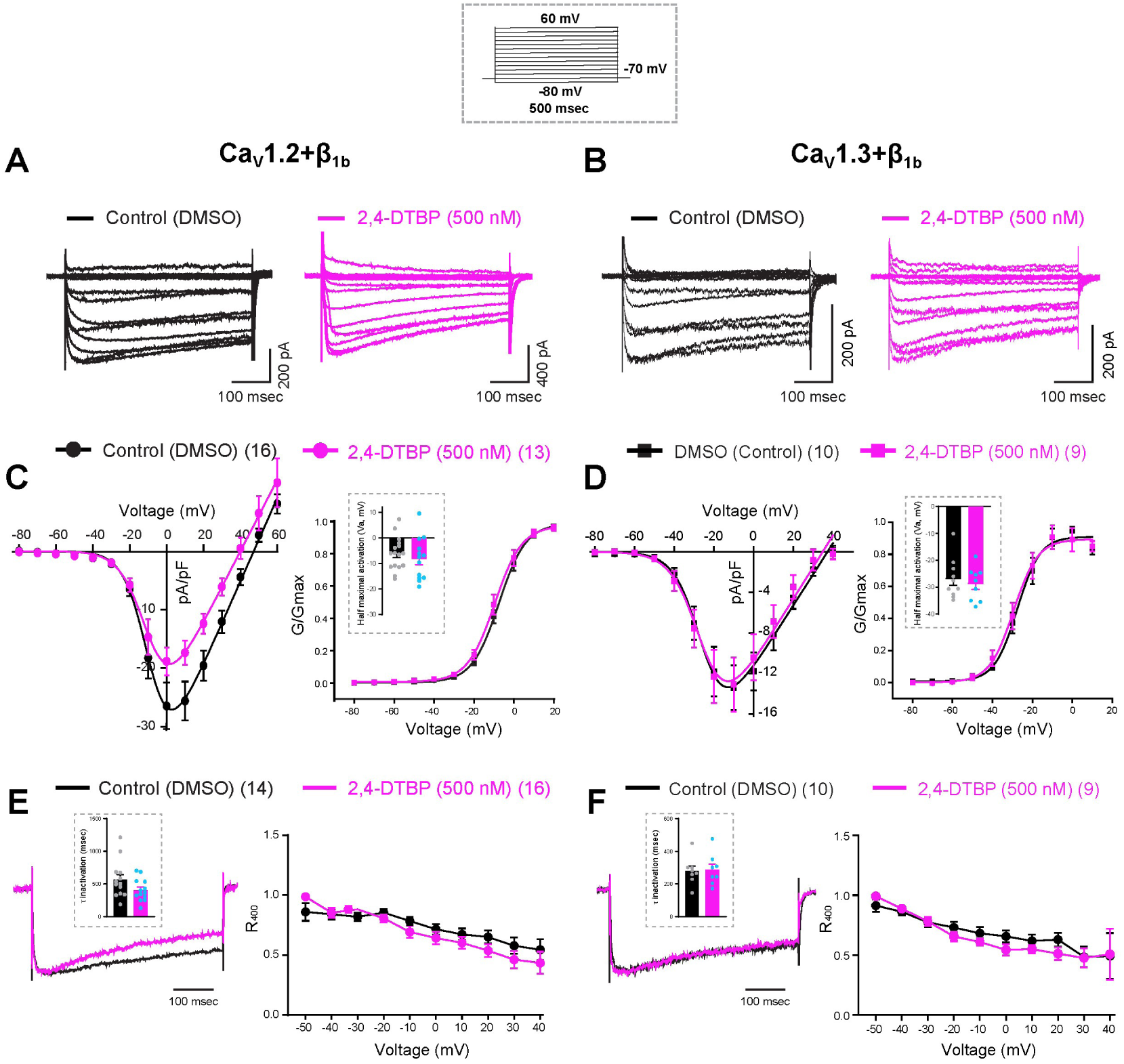
2,4-DTBP does not alter the activation and inactivation properties of Ca_V_1.2 and Ca_V_1.3 channels coexpressed with β_1b_ subunit. **(A and B)** Representative whole-cell current traces of Ca_V_1.2 (**A**) and Ca_V_1.3 (**B**) channels coexpressed with β_1b_ subunit in tsA-201 cells recorded in external solutions containing 500 nM 2,4-DTBP (magenta) or DMSO control (black). The voltage protocol is shown in the dotted box above. **(C and D)** Current density (pA/pF) and normalized conductance (G/G_max_) versus voltage plots of Ca_V_1.2 (**C**) and Ca_V_1.3 (**D**) show no significant differences between 500 nM 2,4-DTBP (magenta) and DMSO control (black). Insets display half-maximal activation voltages. **(E and F)** Normalized current traces of Ca_V_1.2 (**E**) and Ca_V_1.3 (**F**) at 0 mV show channel inactivation with DMSO (black) or 500 nM 2,4-DTBP (magenta). Insets display time constants of inactivation (τ) from exponential fits of 0 mV traces. R_400_ plots represent the ratio of residual current at 400 msec to the peak current amplitude across voltage steps ranging from -50 to +40 mV. All plots represent mean ± SEM. Experimental replicate cell numbers are indicated in brackets. The unpaired Student’s t-test is used for statistical significance. ns, non-significant.

**Figure S9:**
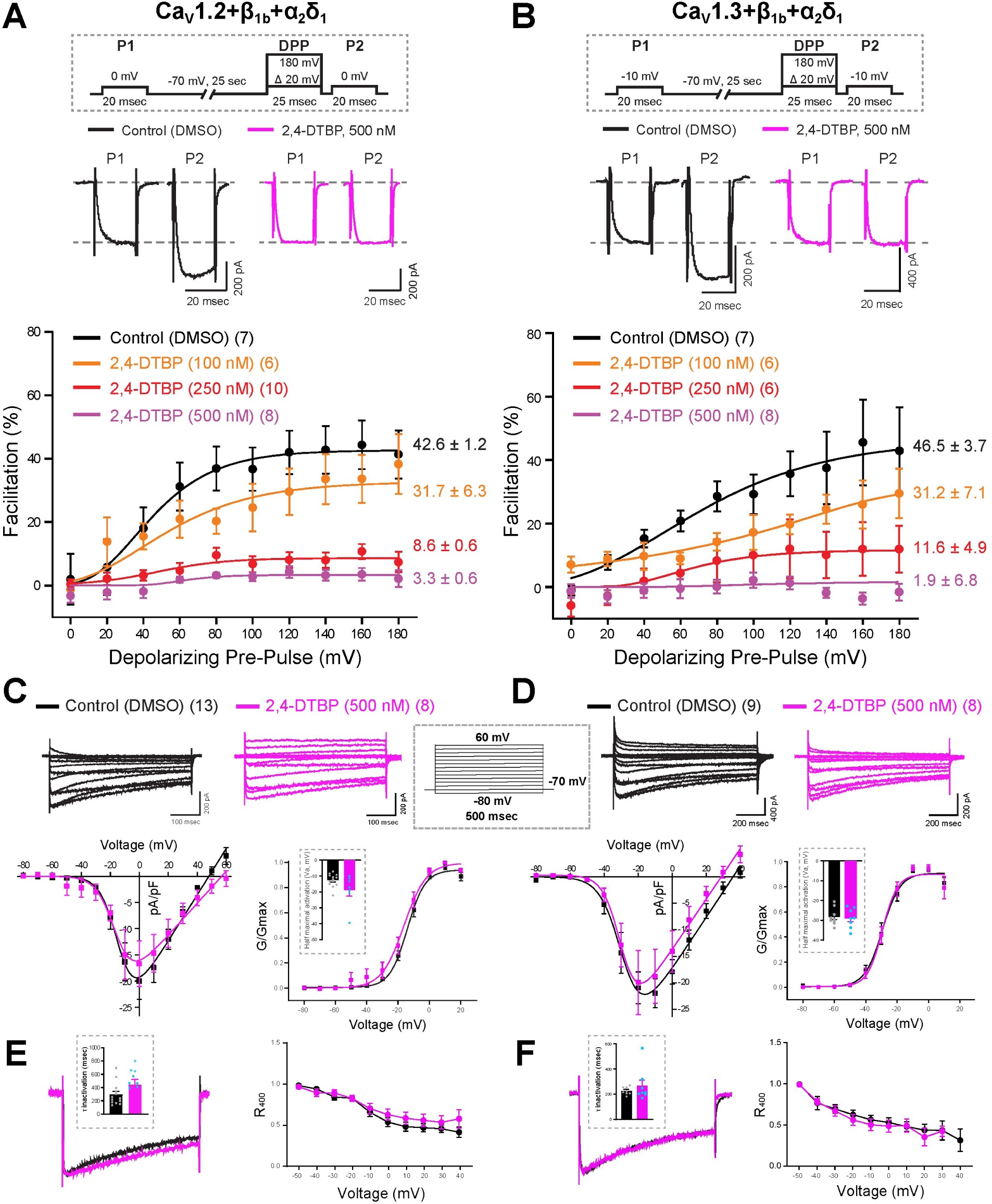
2,4-DTBP potently inhibits the VDF of Ca_V_1.2 and Ca_V_1.3 channels coexpressed with β_1b_ and α_2_δ_1_ subunits, but does not alter their activation and inactivation properties: **(A and B)** Representative whole-cell current traces of L-type Ca_V_1.2 **(A)** and Ca_V_1.3 **(B)** channels coexpressed with β_1b_ and α_2_δ_1_subunits in tsA-201 cells recorded in external solutions containing DMSO or 500 nM 2,4-DTBP before (P1) or after (P2) the DPP to 100 mV. Mean fitted plots (below) show maximum facilitation of Ca_V_1.2 **(A)** and Ca_V_1.3 **(B)** channel currents with DMSO, or 2,4-DTBP at 100, 250, or 500 nM in response to DPP ranging from 0 to 180 mV. Voltage protocol is represented in the dotted box above **(C and D)** Representative whole-cell current traces of Ca_V_1.2 (**C**) and Ca_V_1.3 (**D**) channels coexpressed with β_1b_ and α_2_δ_1_subunits in tsA-201 cells recorded in external solutions containing DMSO (black) or 500 nM 2,4-DTBP (magenta). Voltage protocol is represented in the dotted box above. Current density (pA/pF) and normalized conductance (G/G_max_) voltage plots (below) of Ca_V_1.2 (**C**) and Ca_V_1.3 (**D**) show no significant differences between DMSO control (black) vs 500 nM 2,4-DTBP (magenta). Insets display half-maximal activation voltages. **(E and F)** Normalized current traces of Ca_V_1.2 (**E**) and Ca_V_1.3 (**F**) at 0 mV show channel inactivation with DMSO (black) or 500 nM 2,4-DTBP (magenta). Insets display time constants of inactivation (τ) from exponential fits of 0 mV traces. R_400_ plots represent the ratio of residual current at 400 msec to the peak current amplitude across voltage steps ranging from -50 to +40 mV. All plots represent mean ± SEM. Experimental replicate cell numbers are indicated in brackets. Unpaired Student’s t-test used for statistical significance. ns, non-significant.

**Figure S10:**
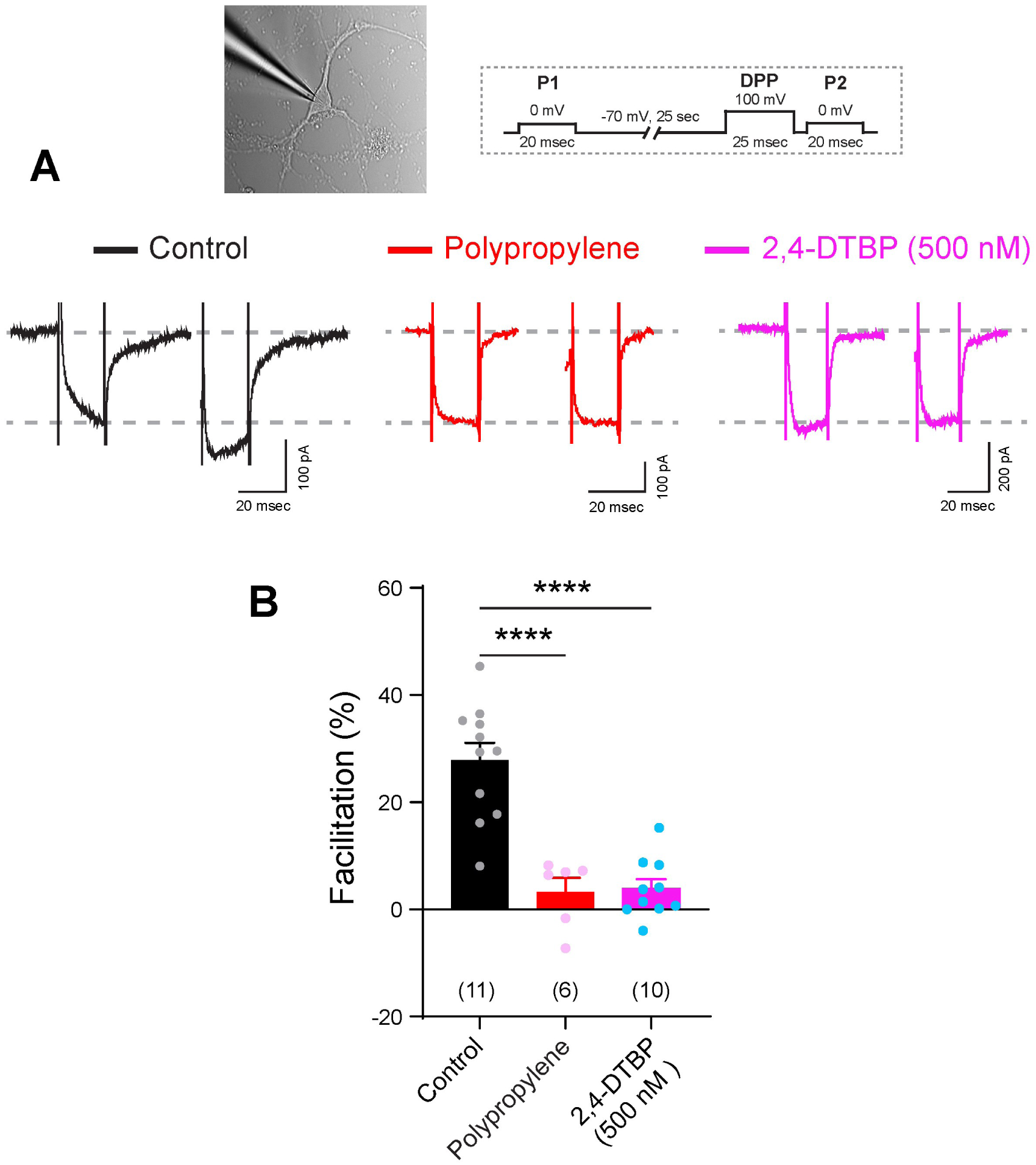
PP-stored solutions and 2,4-DTBP potently inhibit L-type channel VDF in hippocampal pyramidal neurons: **(A)** Representative whole-cell current traces obtained from cultured hippocampal neurons recorded on DIV 7-9 in external solutions stored in glass bottles (black), PP tubes (red), or containing 500 nM 2,4-DTBP (magenta). Whole-cell voltage-clamped hippocampal neuron and voltage protocol shown in the dotted box above. **(B)** Bar plot (Mean ± SEM) quantifications depict the percentage VDF of Ca_V_ channel whole-cell currents, measured as peak current difference between P2 and P1 (0 mV) following DPP to 100 mV. Experimental replicate cell numbers are indicated in brackets. Unpaired Student’s t-test used for statistical significance. ****p ≤ 0.0001.

**Figure S11:**
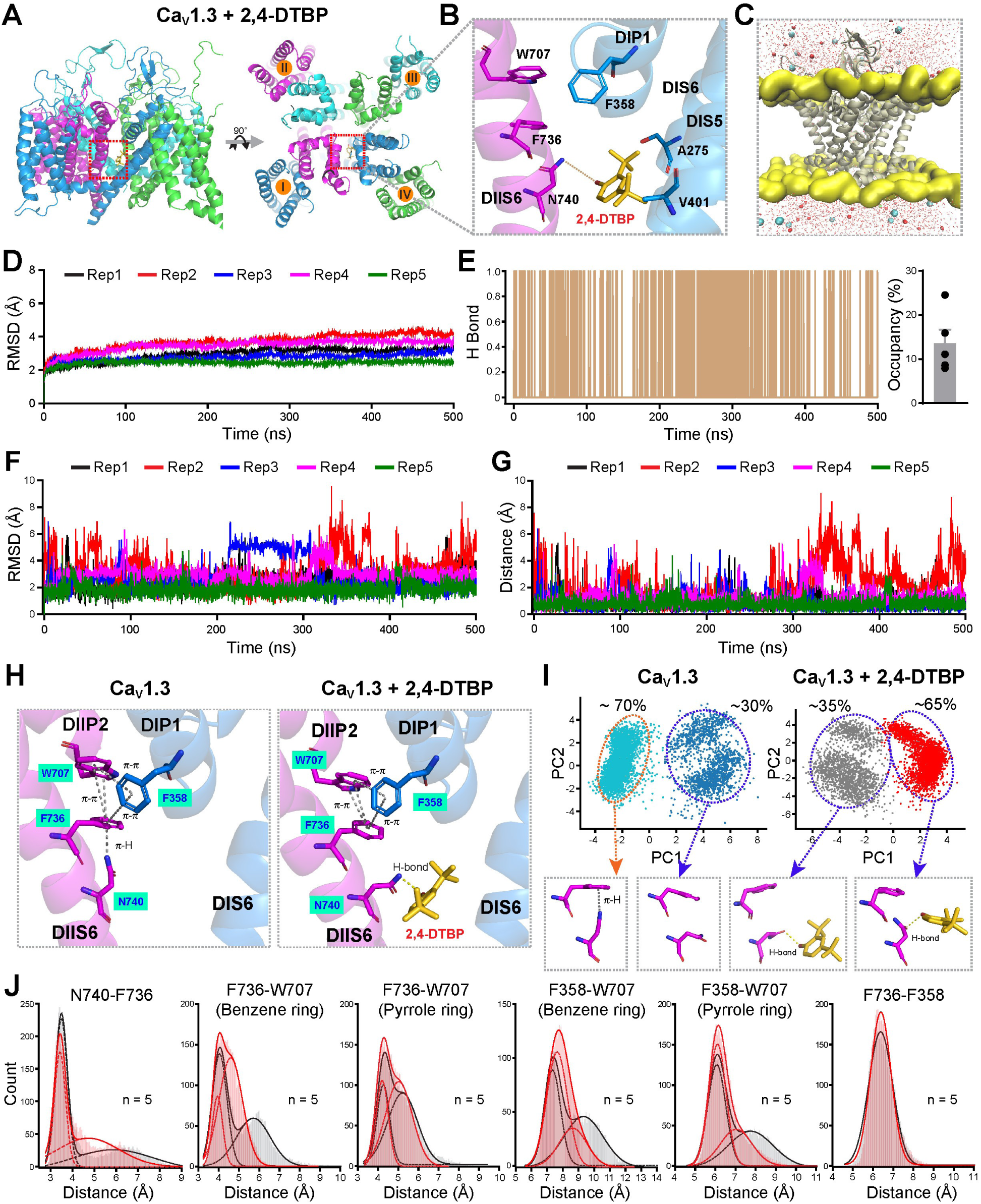
2,4-DTBP disrupts the π-H and π-π quadrangle interaction at the DI and DII PD interface of the Ca_V_1.3 (α_1D_) subunit. **(A)** Side view (left) and top view (right) of the human Ca_V_1.3 protein backbone displaying 2,4-DTBP binding at the DI and DII PD interface. **(B)** Zoomed view of the 2,4-DTBP binding pocket at the DI-DII fenestrations. Key residues forming hydrophobic and H-bonding with 2,4-DTBP in the DIS6, DIS5, and DIIS6 are indicated. **(C)** Graphical representation of the MD simulation setup of the Ca_V_1.3 pore domain with or without 2,4-DTBP. **(D)** Root-mean-square deviation (RMSD) of Ca_V_1.3 backbone during five independent 500 ns MD simulations, showing overall structural stability of the protein-ligand complex. **(E)** Exemplary hydrogen bond occupancy between N740 and 2,4-DTBP during the 500 ns MD-simulation. The bar plot shows the average hydrogen-bond occupancy across five replicates. **(F)** RMSD of 2,4-DTBP in the binding interface during 500 ns MD simulations across five independent replicates, demonstrating ligand stability. **(G)** Distance between the center of mass of 2,4-DTBP and the binding pocket of Ca_V_1.3 in the DI-DII PD interface over the course of 500 ns MD simulations across five independent replicates. **(H)** A zoomed view of the DI and DII PD interface of Ca_V_1.3 reveals feasible π-H and π-π quadrangle interactions among the F358, F736, W707, and N740 residues. **(I)** The PCA of the side chains of F736 and N740 reveals distinct clusters of possible conformations. The representative conformational orientations for each cluster are shown below. **(J)** Centroid-to-centroid distances between the selected residues of Ca_V_1.3 in the absence and presence of 2,4-DTBP over the simulation trajectories.

**Figure S12:**
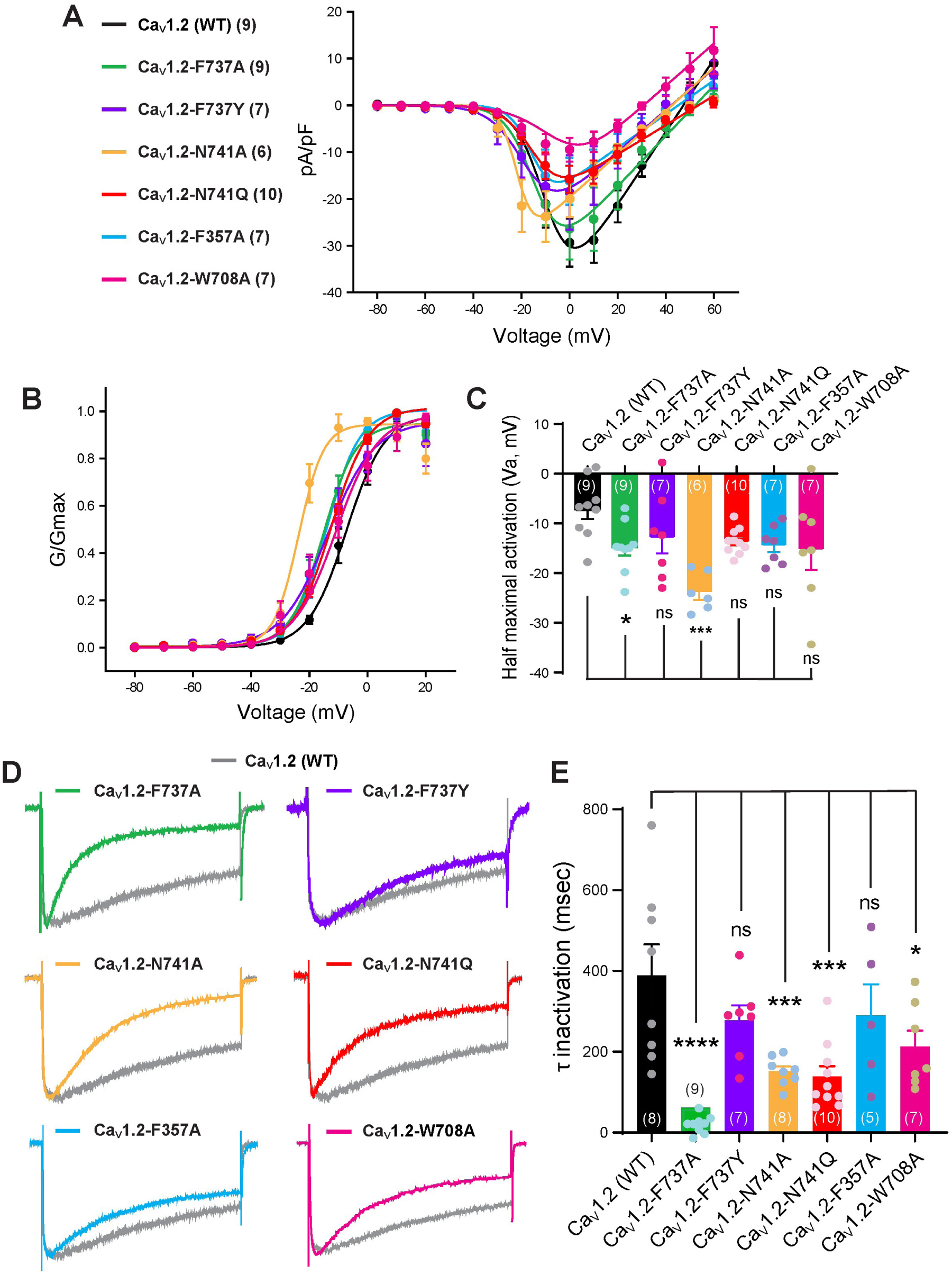
Activation and inactivation kinetics of π-H and π-π quadrangle mutants: **(A)** Current density (pA/pF) vs. voltage plots of Ca_V_1.2 WT and specified mutants. **(B)** Normalized conductance (G/G_max_) vs. voltage plots of Ca_V_1.2 WT and specified mutants. **(C)** Bar plots showing half-maximal activation voltages (V_a_) for Ca_V_1.2 WT and mutants. **(D)** Representative normalized whole-cell current traces of Ca_V_1.2 WT and mutants at 0 mV, displaying inactivation differences. **(E)** Bar plot quantifications of the time constant of inactivation (τ) for Ca_V_1.2 WT and mutants. All plots show mean ± SEM. Experimental replicate cell numbers are indicated in brackets. One-Way ANOVA is used for statistical significance compared to WT. *p ≤ 0.05, **p ≤ 0.01, ***p ≤ 0.001, ****p ≤ 0.0001, ns (non-significant).

**Figure S13:**
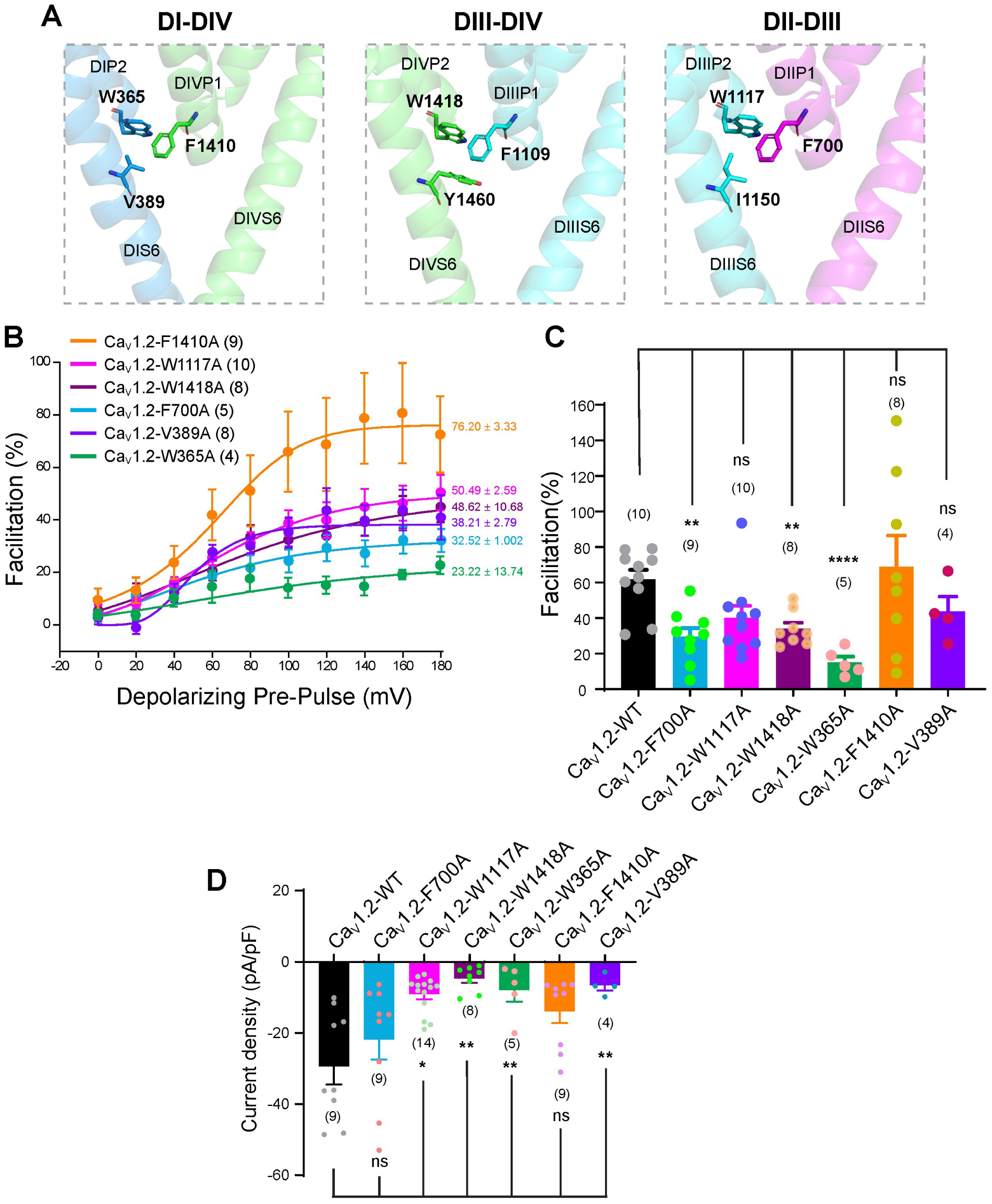
Impact of DII–DIII, DIII–DIV, and DI–DIV PD interface on Ca_V_1.2 VDF: **(A)** Structural representation of DII–DIII, DIII–DIV, and DI–DIV PD interfaces of Ca_V_1.2 showing key aromatic and polar residues. **(B)** Mean fitted plots showing the maximum VDF of specified Ca_V_1.2 mutant channels to DPP, ranging from 0-180 mV. **(C)** Bar plots show VDF quantifications of specified Ca_V_1.2 mutants in comparison with Ca_V_1.2-WT to DPP of 120 mV. **(D)** Bar plots comparing current density of specified Ca_V_1.2 mutant channels vs WT. All plots represent mean ± SEM. Experimental replicate cell numbers are indicated in brackets. One-Way ANOVA is used to assess statistical significance. *p ≤ 0.05, **p ≤ 0.01, ***p ≤ 0.001, ****p ≤ 0.0001, ns, non-significant.

**Figure S14:**
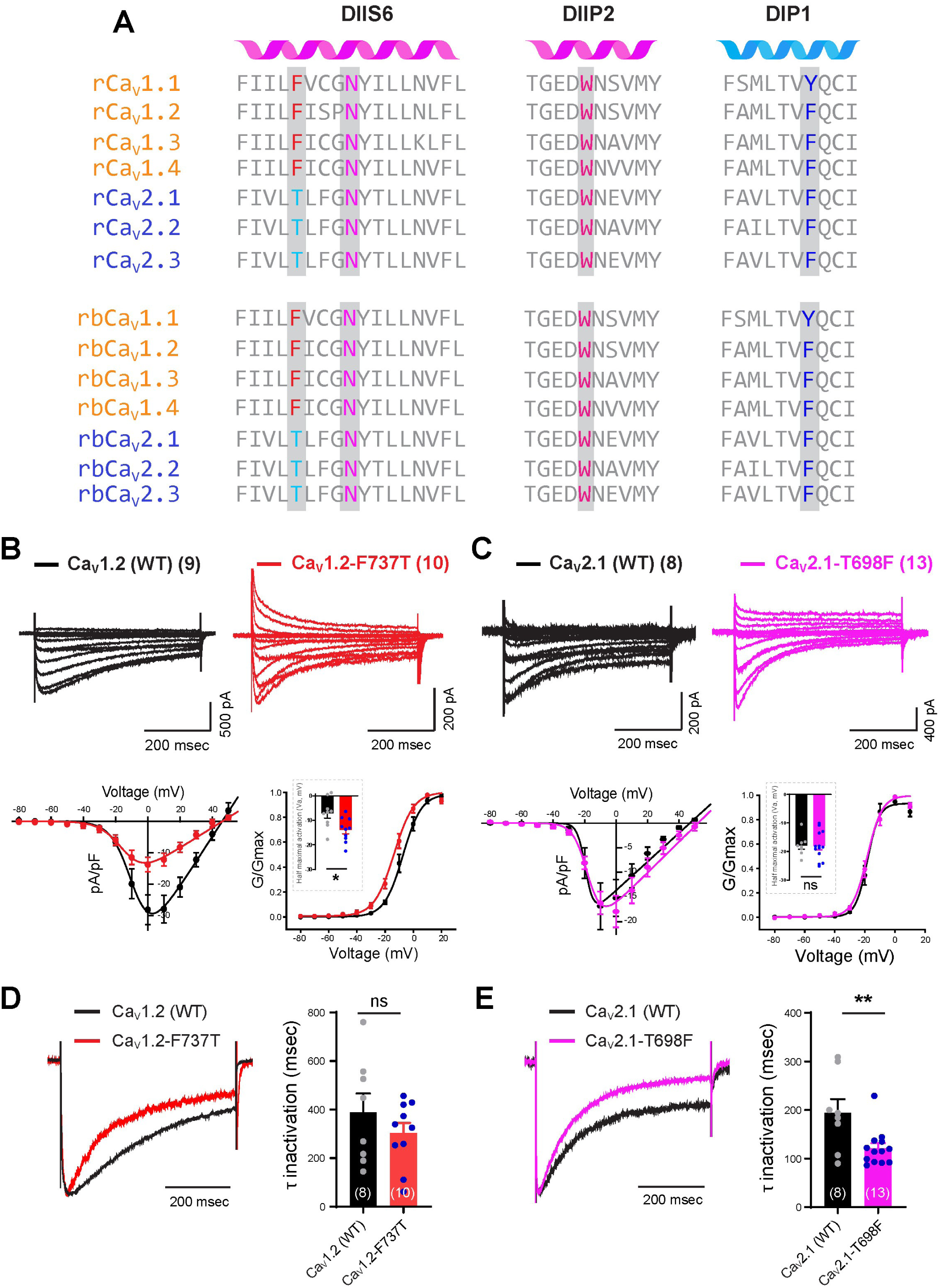
π-H and π-π quadrangle interactions at the DI–DII PD interface in Ca_V_1 and Ca_V_2 HVA channels: **(A)** Aligned amino acid sequences of DIIS6, DIIP2, and DIP1 PD helices from rat (r) and rabbit (rb) L-type (Ca_V_1.1–1.4) and Ca_V_2 (2.1–2.3) channels. Conserved and unique residues participating in the π-H and π-π quadrangle interactions are highlighted. **(B and C)** Representative whole-cell current traces (top) with Current density (pA/pF) and normalized conductance (G/G_max_) vs. voltage plots (below) of WT or mutants of Ca_V_1.2/Ca_V_2.1 channels. The voltage protocol is shown in the dotted box above. Insets display half-maximal activation voltages. **(D and E)** Normalized current traces (left) of WT and mutant Ca_V_1.2/Ca_V_2.1 channels at 0 mV showing channel inactivation. Time constant of inactivation (τ) plots (right) from exponential fits of 0 mV traces. All plots represent mean ± SEM. Experimental replicate cell numbers are indicated in brackets. Unpaired Student’s t-test used for statistical significance. *p ≤ 0.05, **p ≤ 0.01, ns, non-significant.

**Figure S15:**
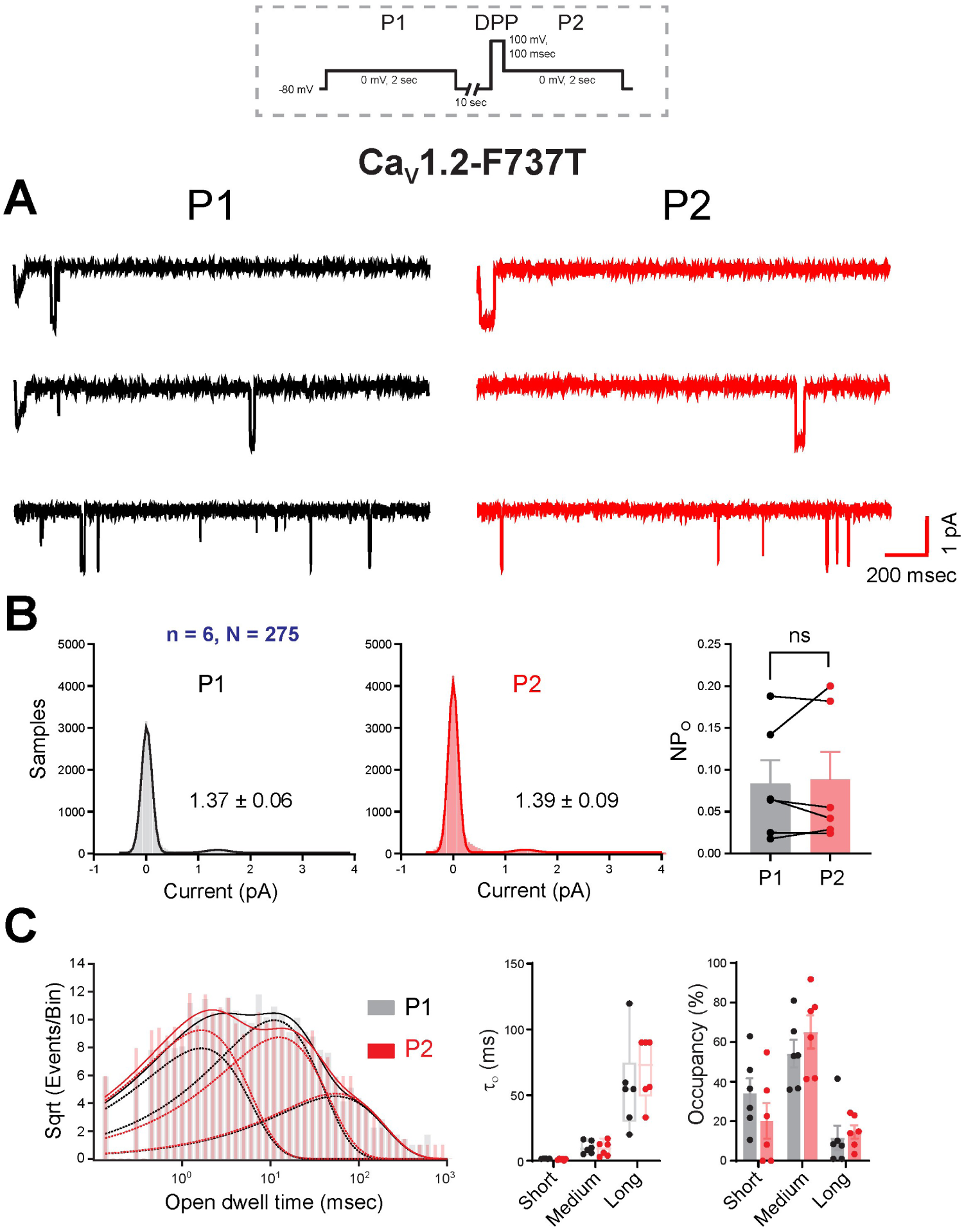
Ca_V_1.2-F737T mutant does not exhibit DPP-induced VDF: **(A)** Representative cell-attached single-channel traces of Ca_V_1.2-F737T mutant coexpressed with β_1b_before and after DPP to 100 mV. The voltage protocol is shown in a dotted box. **(B)** The histogram plots show the single-channel current amplitudes, while the bar plots display the open probability of Ca_V_1.2-F737T cell-attached recordings obtained at 0 mV step of 2 sec before (P1) or after (P2) a DPP of 100 mV for 100 msec. n = number of cells, N = number of single-channel traces. A non-parametric paired Student’s t-test is used to determine statistical significance. **(C)** The logarithmic histogram plot shows the open dwell-time distributions of Ca_V_1.2-F737T channels fitted by nonlinear least squares, while the bar plots display open time constants and their occupancy in short, medium, and long open states. One-way ANOVA with pairwise comparison was used to determine statistical significance. All plots represent mean ± SEM. ns, p > 0.05 (non-significant).

---

**Table S1:** VDF, activation and inactivation parameters of WD and point-mutated Ca_V_1.2 and Ca_V_1.3, and Ca_V_2.1 channels.

| Sl. No | Channel | VDF at 120 mV | Current Density (pA/pF) | V<sub>a</sub> (mV) | E<sub>rev</sub> (mV) | Tau Inactivation (msec) | Cell No. (n) |
| --- | --- | --- | --- | --- | --- | --- | --- |
| 1 | Cav1.2+β<sub>1</sub>b | 61.69 ± 5.43 | -29.37 ± 5.11 | -7.13 ± 1.98 | 49.01 ± 2.50 | 388.9 ± 76.95 | 9 |
| 2 | Cav1.3+ β<sub>1</sub>b | 41.32 ± 4.65 | -12.23 ± 2.44 | -26.63 ± 3.65 | 37.71 ± 2.93 | 371.6 ± 81.57 | 8 |
| 3 | Cav1.2+β<sub>1</sub>b+α<sub>2</sub>δ<sub>1</sub> | 36.64 ± 5.75 | -44.30 ± 6.18 | -13.50 ± 0.64 | 49.80 ± 1.79 | 129.55 ± 16.04 | 9 |
| 4 | Cav1.3+β<sub>1</sub>b+α<sub>2</sub>δ<sub>1</sub> | 43.42 ± 7.72 | -27.59 ± 6.16 | -27.59 ± 1.20 | 43.36 ± 1.24 | 231.83 ± 36.68 | 6 |
| 5 | Cav1.2+β<sub>1</sub>b (PP) | 3.97 ± 1.87 (****) | -27.27 ± 3.38 | -9.67 ± 1.77 | 48.84 ± 1.67 | 322.4 ± 120.94 | 12 |
| 6 | Cav1.3+β<sub>1</sub>b (PP) | 3.09 ± 2.57 (****) | -11.25 ± 1.59 | -26.92 ± 1.36 | 47.47 ± 2.71 | 425.55 ± 101.1 | 9 |
| 7 | Cav1.2+ β<sub>1</sub>b+α<sub>2</sub>δ<sub>1</sub> (PP) | 2.67 ± 2.71 (****) | -44.38 ± 7.95 | -16.09 ± 1.25 | 47.98 ± 1.18 | 110.67 ± 12.77 | 9 |
| 8 | Cav1.3+β<sub>1</sub>b+α<sub>2</sub>δ<sub>1</sub> (PP) | -0.23 ± 2.99 (***) | -24.76 ± 5.67 | -29.75 ± 1.71 | 39.41 ± 1.67 | 185.14 ± 17.20 | 7 |
| 9 | Cav1.2+β<sub>1</sub>b (DMSO) | 55.76 ± 6.99 | -26.46 ± 3.97 | -6.16 ± 1.60 | 47.49 ± 1.35 | 565.08 ± 78.02 (ns) | 16 |
| 10 | Cav1.3+β<sub>1</sub>b (DMSO) | 35.46 ± 6.12 | -13.39 ± 2.20 | -26.97 ± 2.36 | 43.33 ± 2.34 | 279.12 ± 30.25 | 10 |
| 11 | Cav1.2+β<sub>1</sub>b+α<sub>2</sub>δ<sub>1</sub> (DMSO) | 42.0 ± 6.83 | -20.01 ± 3.50 | -12.6 ± 1.12 | 50.47 ± 1.48 | 295.30 ± 46.7 | 13 |
| 12 | Cav1.3+β<sub>1</sub>b+α<sub>2</sub>δ<sub>1</sub> (DMSO) | 40.77 ± 8.11 | -21.97 ± 2.69 | -28.23 ± 1.42 | 40.62 ± 2.67 | 227.25 ± 10.58 | 9 |
| 13 | Cav1.2+ β<sub>1</sub>b (2,4-DTBP-500 nM) | 3.90 ± 1.77 (****) | -18.86 ± 2.41 | -8.2 ± 2.28 | 43.10 ± 2.54 | 405.23 ± 47.21 | 13 |
| 14 | Cav1.3+β<sub>1</sub>b (2,4-DTBP-500 nM) | 2.18 ± 1.51 (***) | -13.09 ± 2.69 | -28.77 ± 2.08 | 42.56 ± 5.45 | 287.75 ± 33.95 | 9 |
| 15 | Cav1.2+β<sub>1</sub>b+α<sub>2</sub>δ<sub>1</sub> (2,4-DTBP-500 nM) | 3.75 ± 1.39 (****) | -16.68 ± 4.33 | -18.74 ± 3.76 | 57.78 ± 6.52 | 442.0 ± 80.36 | 7 |
| 16 | Cav1.3+β<sub>1</sub>b+α<sub>2</sub>δ<sub>1</sub> (2,4-DTBP-500 nM) | 2.95 ± 2.03 (***) | -19.08 ± 5.10 | -28.82 ± 1.74 | 32.50 ± 3.91 | 266.25 ± 45.47 | 8 |
| 17 | Cav1.2-F737A | 1.72 ± 4.9 (****) | -26.42 ± 6.59 | -14.82 ± 1.67 | 52.96 ± 2.35 | 61.44 ± 7.24 (***) | 9 |
| 18 | Cav1.2-F737T | 2.99 ± 2.24 (****) | -13.73 ± 2.29 (*) | -13.9 ± 1.56 | 44.49 ± 2.95 | 302.5 ± 42.11 | 10 |
| 19 | Cav1.2-F737Y | 59.37 ± 10.72 (ns) | -18.6 ± 7.94 | -12.68 ± 3.37 | 44.53 ± 4.26 | 278.3 ± 36.62 | 7 |
| 20 | Cav1.2-N741A | $-0.78 \pm 3.16$<br>(****) | $-19.99 \pm 3.90$ | $-23.75 \pm 1.60$ | $51.57 \pm 2.96$ | $151.5 \pm 12.32$ | 6 |
| 21 | Cav1.2-N741Q | $1.85 \pm 5.24$<br>(****) | $-15.80 \pm 2.88$ | $-13.58 \pm 0.82$ | $56.33 \pm 2.59$ | $138.4 \pm 26.06$ | 10 |
| 22 | Cav1.2-F357A | $21.85 \pm 5.60$<br>(***) | $-16.21 \pm 4.99$ | $-14.32 \pm 1.46$ | $46.28 \pm 2.72$ | $289.8 \pm 77.50$ | 7 |
| 23 | Cav1.2-W708A | $12.45 \pm 8.75$<br>(***) | $-9.57 \pm 2.54$<br>(**) | $-15.18 \pm 4.25$ | $32.80 \pm 2.43$ | $212.9 \pm 39.50$ | 7 |
| 24 | Cav1.2-F700A | $29.46 \pm 4.97$<br>(***) | $-21.88 \pm 5.60$ | $-20.14 \pm 2.41$ | $44.32 \pm 2.95$ | $362.75 \pm 79.46$ (ns) | 9 |
| 25 | Cav1.2-F1410A | $68.79 \pm 17.67$<br>(ns) | $-13.93 \pm 3.29$<br>(*) | $-8.55 \pm 2.14$ | $48.02 \pm 3.02$ | $364.44 \pm 58.90$ (ns) | 9 |
| 26 | Cav1.2-F1109A | -- | No Expression | No Expression | -- | -- | -- |
| 27 | Cav1.2-Y1460A | -- | No Expression | No Expression | -- | -- | -- |
| 28 | Cav1.2-V389A | $43.61 \pm 8.46$ | $-6.6 \pm 1.42$ (*) | $-4.51 \pm 1.96$ | $59.12 \pm 5.84$ | | 4 |
| 29 | Cav1.2-W1117A | $40.16 \pm 6.78$<br>(*) | $-9.13 \pm 1.36$<br>(***) | $2.03 \pm 3.35$ | $31.89 \pm 2.46$ | -- | 14 |
| 30 | Cav1.2-W1418 | $33.94 \pm 3.44$<br>(***) | $-4.68 \pm 1.254$<br>(***) | $7.47 \pm 3.50$ | $40.13 \pm 4.05$ | | 8 |
| 31 | Cav1.2-W365A | $15.10 \pm 3.22$<br>(****) | $-5.41 \pm 2.80$<br>(**) | $0.07 \pm 4.77$ | $49.48 \pm 2.57$ | -- | 5 |
| 32 | Cav2.1+β<sub>1b</sub> | $2.75 \pm 1.17$ | $-15.21 \pm 3.69$ | $-17.87 \pm 1.29$ | $47.01 \pm 1.08$ | $194.5 \pm 27.89$ | 8 |
| 33 | Cav2.1-T698F | $40.14 \pm 7.41$<br>(****) | $-17.95 \pm 3.21$ | $-18.02 \pm 1.38$ | $52.55 \pm 2.88$ | $121.3 \pm 10.42$ | 13 |

PP: Polypropylene; DMSO: Dimethyl Sulfoxide; 2,4-DTBP: 2,4-Ditert-butylphenol; WT: Wild type. All values represent mean ± SEM. \* $p \leq 0.05$ , \*\* $p \leq 0.01$ , \*\*\* $p \leq 0.001$ , \*\*\*\* $p < 0.0001$ . Cav1.2 or Cav1.3 channels coexpressed with $\alpha_2\delta_1$ are compared with respective channels coexpressed with $\beta_{1b}$ . Point mutated Cav1.2 and Cav2.1 channels are compared with respective WT channels.

**Table S2:**
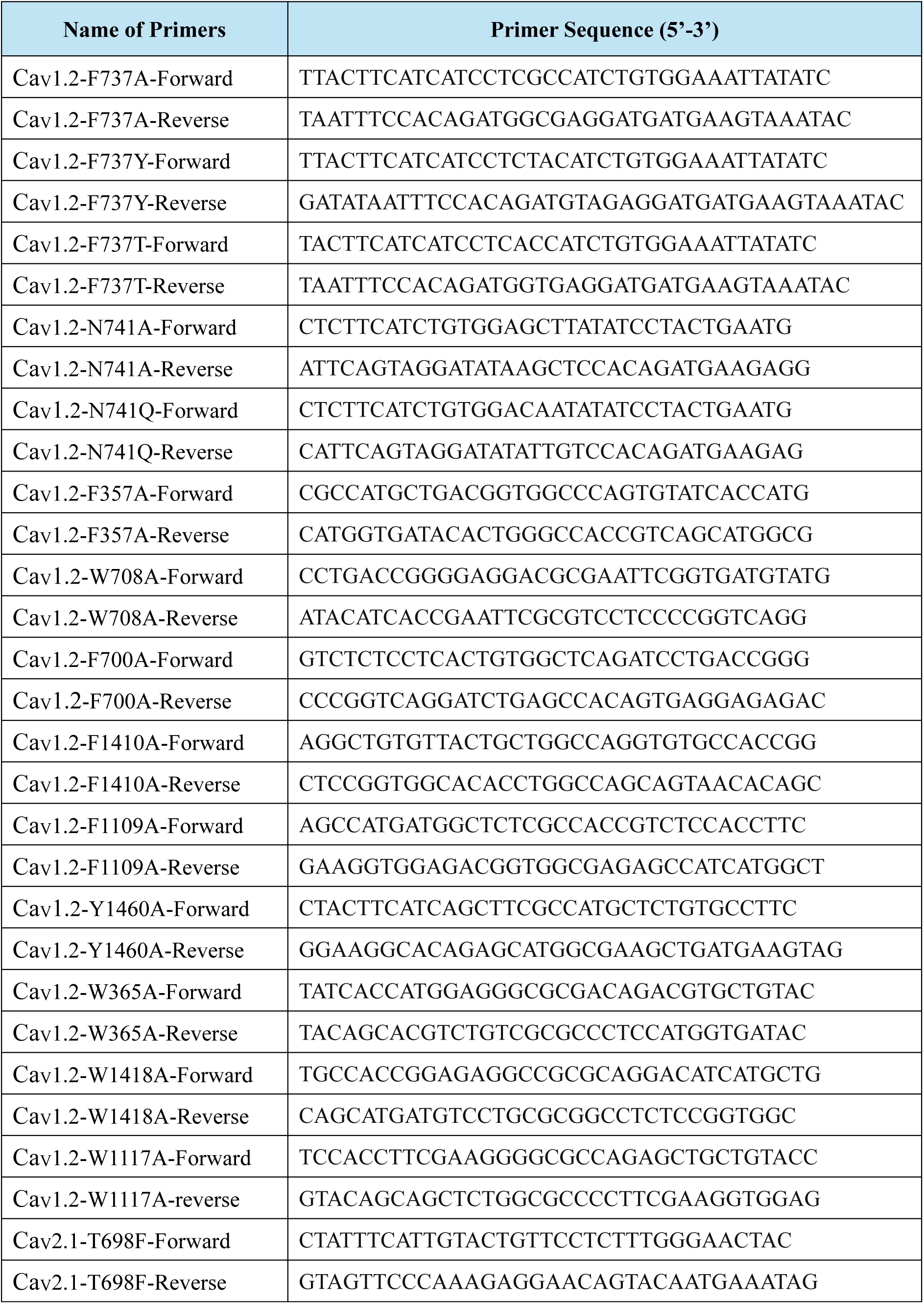
Lists of point mutagenesis primers used.

**Table S3:** List of accession numbers used for multiple sequence alignment.

| **Sl. No** | **Channel Name** | **UniProt id** | **Organism** |
| --- | --- | --- | --- |
| 1 | Cav1.1 | Q13698 | Human |
| 2 | Cav1.2 | Q13936 | Human |
| 3 | Cav1.3 | Q01668 | Human |
| 4 | Cav1.4 | O60840 | Human |
| 5 | Cav2.1 | O00555 | Human |
| 6 | Cav2.2 | Q00975 | Human |
| 7 | Cav2.3 | Q15878 | Human |
| 8 | Cav1.1 | Q02789 | Mouse |
| 9 | Cav1.2 | Q01815 | Mouse |
| 10 | Cav1.3 | Q99246 | Mouse |
| 11 | Cav1.4 | Q9JIS7 | Mouse |
| 12 | Cav2.1 | P97445 | Mouse |
| 13 | Cav2.2 | O55017 | Mouse |
| 14 | Cav2.3 | Q61290 | Mouse |
| 15 | Cav1.1 | Q02485 | Rat |
| 16 | Cav1.2 | P22002 | Rat |
| 17 | Cav1.3 | P27732 | Rat |
| 18 | Cav1.4 | Q923Z7 | Rat |
| 19 | Cav2.1 | P54282 | Rat |
| 20 | Cav2.2 | Q02294 | Rat |
| 21 | Cav2.3 | Q07652 | Rat |
| 22 | Cav1.1 | P07293 | Rabbit |
| 23 | Cav1.2 | A0A5F9CX63 | Rabbit |
| 24 | Cav1.3 | G1T5I1 | Rabbit |
| 25 | Cav1.4 | G1SNI0 | Rabbit |
| 26 | Cav2.1 | P27884 | Rabbit |
| 27 | Cav2.2 | Q05152 | Rabbit |
| 28 | Cav2.3 | Q02343 | Rabbit |

## Notes

### Competing Interest Statement

The authors have declared no competing interest.

### Summary of Updates

New experimental data are added to the manuscript

